# Statistical Inference of a Convergent Antibody Repertoire Response to Influenza Vaccine

**DOI:** 10.1101/025098

**Authors:** Nicolas Strauli, Ryan Hernandez

## Abstract

**Background:** Vaccines dramatically affect an individual’s adaptive immune system, and thus provide an excellent means to study human immunity. Upon vaccination, the B cells that express antibodies (Abs) that happen to bind the vaccine are stimulated to proliferate and undergo mutagenesis at their Ab locus. This process may alter the composition of B cell lineages within an individual, which are known collectively as the antibody repertoire (AbR). Antibodies are also highly expressed in whole blood, potentially enabling unbiased RNA sequencing technologies to query this diversity. Less is known about the diversity of AbR responses across individuals to a given vaccine and if individuals tend to yield a similar response to the same antigenic stimulus.

**Methods:** Here we implement a bioinformatic pipeline that extracts the AbR information from a time-series RNA-seq dataset of 5 patients who were administered a seasonal trivalent influenza vaccine (TIV). We harness the detailed time-series nature of this dataset and use methods based in functional data analysis (FDA) to identify the B cell lineages that respond to the vaccine. We then design and implement rigorous statistical tests in order to ask whether or not these patients exhibit a convergent AbR response to the same TIV.

**Results:** We find that high-resolution time-series data can be used to help identify the Ab lineages that respond to an antigenic stimulus, and that this response can exhibit a convergent nature across patients inoculated with the same vaccine. However, correlations in AbR diversity among individuals prior to inoculation can confound inference of a convergent signal unless it is taken into account.

**Conclusions:** We developed a framework to identify the elements of an AbR that respond to an antigen. This information could be used to understand the diversity of different immune responses in different individuals, as well as to gauge the effectiveness of the immune response to a given stimulus within an individual. We also present a framework for testing a convergent hypothesis between AbRs; a hypothesis that is more difficult to test than previously appreciated. Our discovery of a convergent signal suggests that similar epitopes do select for antibodies with similar sequence characteristics.

## Background

Since the administration of the first designed vaccine by Edward Jenner in 1796 [1], vaccines have proven indispensible for both medicine and medical research. Vaccines are among the rare achievements of science that have fundamentally changed modern life. Perhaps less well known, vaccines also provide a standardized, safe, and ethical way to directly study human adaptive immunity [2]. Most vaccines confer resistance to a given pathogen by stimulating the patient’s population of B cells to produce antibodies (Abs) against the inoculated antigens. Each clonal lineage of B cells express a different Ab, and the conglomerate of B cells within an individual make up their antibody repertoire (AbR).

Interestingly, the process by which Abs are adapted to more specifically target an insulting antigen is essentially identical to evolution by natural selection. To wit, during B cell development a vast amount of genetic diversity is generated by a series of somatic mutagenic steps, after which, variants that are able to bind an antigen strongly will be positively selected to proliferate [3,4]. The first diversity-generating step in B cell development is a process of somatic recombination that takes place in the bone marrow. The mature Ab protein is composed of two identical light chains and two identical heavy chains. A light chain can be of either the lambda (IGL) or kappa (IGK) variety, whereas the heavy chain has only one possibility (IGH), and the loci encoding these three chains reside in distinct regions of the genome. Here, the Variable (V), Diversity (D), and Joining (J) gene segments in the IGH locus, and V and J gene segments in the light chain loci will recombine [5–7]. Diversity is generated both by selecting one combination out of all the possible combinations of V, (D) and J genes, as well as by the random insertion and deletion of genetic information at the junctions of these gene segments [8]. Further, once a mature B cell binds an antigen, it will be recruited to a lymph follicle and enter a structure known as the germinal center where a process of somatic hyper-mutation (SHM) takes place [3,4]. Random point mutations are smattered onto the variable region of the Ab locus—the area that is responsible for binding antigen—and if these mutations result in high binding affinity, that B cell clone will receive signals to proliferate. This process generates lineages of B cells specific for a given antigen. These mutagenic steps together result in a high concentration of mutations occurring in a region of the Ab called the complementary determining region 3 (CDR3), which happens to be the region of the Ab that tends to physically interact with antigen. Because of this, the sequence encoding the CDR3 is often used for clonal analysis of Abs, where Abs with the same CDR3 sequence are assumed to be clones. The net effect of this evolutionary process produces extreme temporal dynamism within the AbR, as different lineages grow and shrink in response to different antigenic stimuli [9].

Advancements in next-generation sequencing (NGS) methods have led to recent work in characterizing the AbR’s response to a variety of stimuli [9–21] (see Galson *et al* [2] for a review). However, most of this work has focused on methods development, and there has been comparatively little work focusing on what can actually be learned from these data. Contrary to this trend, Greiff *et al.* [22] recently employed a machine learning approach to classify patients’ immune status using their AbR sequence data. Much work remains to be done in this relatively new area of research. For example, the overall changes in a patient’s AbR could be used to quantitatively assess the response to vaccination. Of particular interest is the ability to use changes in expression of individual Abs to identify which specific monoclonal Abs (mAbs) bind a given antigen. To address this gap in knowledge, we here seek to leverage time-series information of five patients’ AbRs in order to infer the elements that are responding to a trivalent influenza vaccine (TIV).

A particularly useful and intuitive way to model time-series data is to use methods within the greater discipline of functional data analysis (FDA) [23,24]. As opposed to multivariate data analysis (MDA)—which treats each datum as a finite dimensional vector of observations—FDA treats each datum as a continuous function over some dimension, which is often (as in our case) time. FDA-based methods have a rich history of being used for identifying differentially expressed genes over time [25–28], and have the advantage of easily incorporating uneven time-point sampling, and measurement error into each gene’s functional model. FDA is also an intuitive way to model gene expression, as each gene’s expression level in a tissue is indeed continuously fluctuating over continuous time. Here, we use an FDA-based method presented by Wu and Wu [27], and apply it to time-series AbR data to identify the components of patients’ AbR that are responding to a standard TIV.

There is a plethora of time-series gene expression data that have been used to identify genes involved in pathogen defense [29], autoimmunity [29], and vaccine response [30,31]. The longitudinal and cross-sectional nature of these studies allowed the authors to identify the genes that were consistently differentially expressed in response to the given antigenic stimulus across patients. One could perform a similar analysis using a time-series AbR dataset to help identify the determinants of immunity. However, with the exception of Liao *et al.* [32] and Laserson *et al.* [9], few detailed time-series datasets on the AbR exist [9,32]. If RNA-seq were performed on an antibody expressing tissue (for example peripheral blood mononuclear cells, or PBMCs), theoretically, many of the RNA transcripts in the data would originate from Ab loci. Should this be the case, there will exist much AbR information within the data that simply need to be bioinformatically mined out. This approach has been used in the context of cancer research to both identify the Ab sequence of the cancerous B cell lineage in Chronic Lymphocytic Leukemia patients [33], as well as to characterize AbR diversity in solid tumor samples [34]. In this study, we developed and implemented such a pipeline on the Henn *et al.* 2013 [31] transcriptomic dataset in order to probe the AbR’s response to a standard TIV.

There have been several reports of convergent evolutionary signals between independent AbRs that were exposed to a similar antigenic stimulus. This has been shown for dengue virus infections [21], broadly neutralizing Abs against Human Immunodeficiency Virus [15,18], and influenza vaccination [35,36]. With the exception of Parameswaran *et al.* [21], these studies relied largely on qualitative evidence for convergence, where Ab sequences from independent patients either cluster closely together on a dendogram [15], or have strikingly similar sequence and/or structural characteristics [18,35,36]. While these examples of AbR convergence may be intuitively convincing, few methods have been developed to statistically test for a convergent AbR response across patients. The importance of statistical analyses can be illustrated by the high correlation of Ab gene expression in different individuals [37,38]. That is, if an Ab gene is expressed highly in one individual, it will tend to also be highly expressed in another individual. In order to soundly establish a convergent signal between patients’ AbRs, this correlation in background gene expression must first be taken into account. To resolve this, we developed and implemented a statistical methodology that incorporates the baseline similarity between individual AbRs when testing for a convergent signal.

In this study, we first present a bioinformatic pipeline for extracting AbR information from RNA-seq data. We then go on to use FDA-based methods to characterize the Ab response of several patients to a standard TIV. Finally, we present and implement statistical tests for a convergent Ab response between patients to the same TIV. We find that a detailed time-series dataset can be used to identify Abs that are putatively binding a vaccine, and that—after controlling for background AbR similarities—these vaccine targeting Abs can exhibit similar sequence characteristics across patients.

## Methods

### Data creation

The RNA-seq dataset for this study was generated by Henn *et al.* [31] The experimental design was as follows: 5 patients were vaccinated with the 2010 seasonal TIV, and blood was drawn from each patient for 11 days, from day 0 (the day of the vaccination) to day 10 post vaccination. Each patient/time-point sample was divided into 2 sample-types: PBMCs, and sorted B cells. RNA-seq was performed on both the PBMC and B cell sample-types from each time point for all patients. Importantly, the two different sample-types from each sample provide relatively independent technical replicates to gauge the accuracy of our bioinformatic pipeline, described below.

For a detailed description of sample processing and RNA sequencing see [31]. Briefly, PBMCs were isolated using a discontinuous Ficoll gradient centrifugation, and B cells were enriched from heparinized whole blood with RosetteSep Immunodensity separation (Stemcell Technologies, Vancouver, BC, Canada). RNA was extracted with the Qiagen RNeasy micro kit. Barcodes were assigned to each patient/time-point/sample-type, and sequencing libraries were prepared with Illumina TruSeq RNA kits as recommended by Illumina, using 100 ng total RNA as input. The read length was 65 bases, and the mean read depth across patient/time-point/sample-types was 13,724,354.04 reads, with a range of 8,262,317-17,777,695 reads.

### Computational pipeline

State-of-the-art tools for aligning RNA-seq reads to a reference genome, such as TopHat2 [39], were not designed, and are ill-equipped, to handle the various eccentricities of Ab RNA (such as VDJ gene segment recombination, as well as the high number of mutations expected from both VDJ recombination and SHM). Similar to others [33,34], we therefore developed a bioinformatic pipeline that will harvest the Ab transcripts buried in the multitude of reads from an RNA-seq dataset. All code for running this bioinformatic pipeline is available upon request. Conceptually, the pipeline consists of a negative selection step to weed out all non-Ab encoding transcripts, followed by a positive selection step to identify Ab encoding reads (Fig. 1A). For the negative selection step, we first created a whole genome reference sequence where all Ab encoding loci in the genome were masked out. We then used TopHat2 to map all reads to this masked-reference genome. Reads that successfully mapped to the masked genome were discarded. We hypothesized that some fraction of the unmapped reads are true Ab sequences. To identify them, we used IgBLAST [40] to positively select for Ab encoding transcripts. We used a stringent threshold (e-value ≤ 10^−20^) to select the best matching germline Ab gene (including V, D, and J genes) and the CDR3 sequence (if available) from the IgBLAST output.

**Figure 1.**
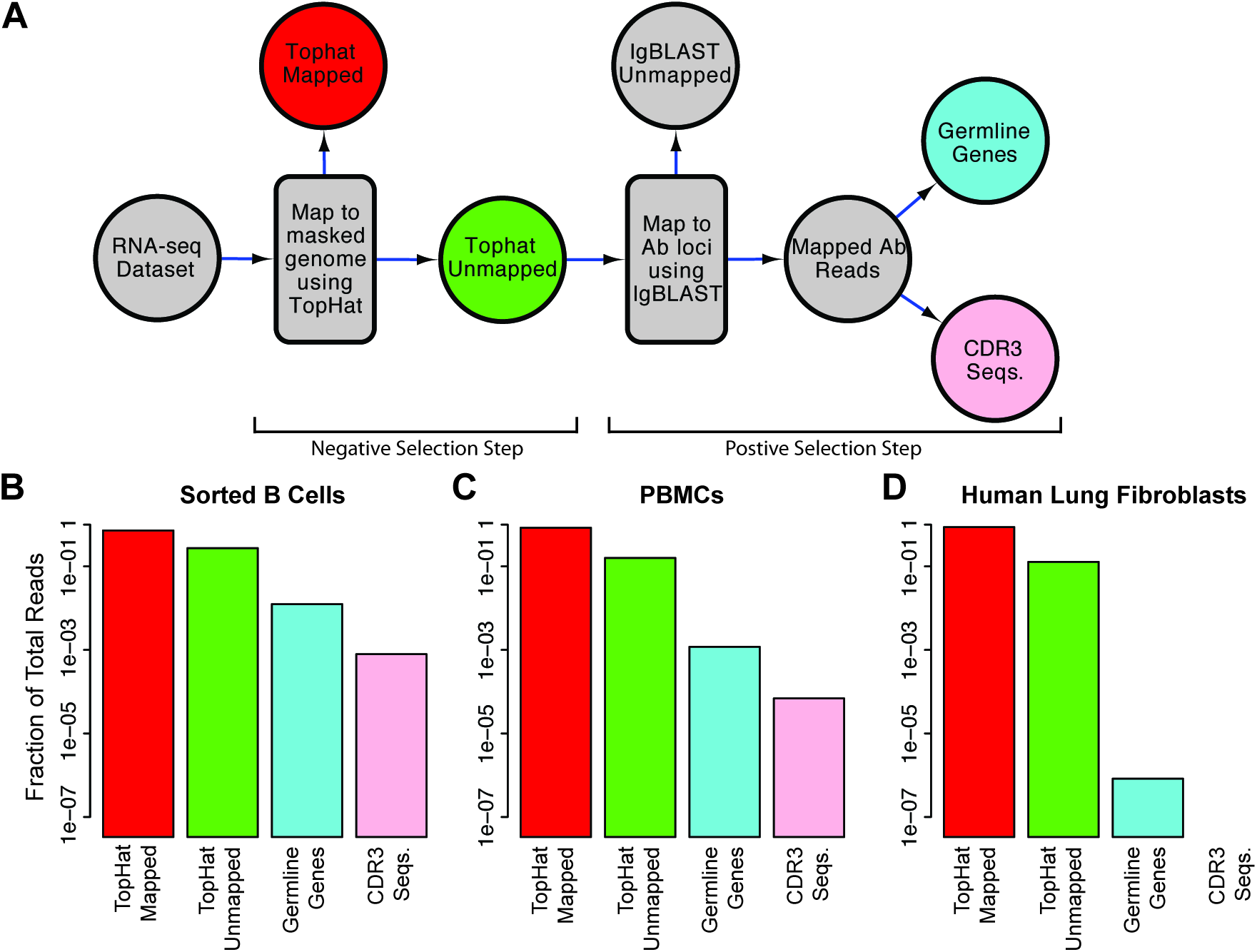
Bioinformatic pipeline. (A) Flow diagram of the steps in our bioinformatic pipeline for harvesting Ab reads from a RNA-seq dataset. The pipeline consists of a negative selection step using TopHat2 [39] where non-Ab reads are mapped to a masked reference genome, followed by a positive selection step using IgBLAST [40] where Ab reads are mapped to reference germline Ab sequences. (B) Fraction of reads retrieved for certain steps in the pipeline, in 3 different tissues, out of the number of TopHat mapped reads (red). The colors of the bars correspond to the colors of the steps in (A).

### Overall Ab abundance and Ab gene expression

We first would like to estimate the overall abundance (or expression) of Abs in each sample. Let *T* be the total number of days in the study, with *t* ∈ [0,*T*], and *P* be the total number of patients, with *i* ∈ [1,*P*]. For a given patient *i*, and timepoint *t*, if *M*_*i,t*_ is the total number of reads that map to an Ab locus with an e-value ≤ 10^−20^, and *N*_*i,t*_ is the total number of reads that map to anything in the Ab-masked genome (the red circle in Fig. 1A), then Ab abundance, *A*_*i,t*_, for that patient/time-point can be calculated as,

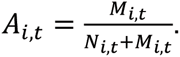

Because *M*_*i,t*_ was very small relative to *N*_*i,t*_, we approximated Ab abundance as,

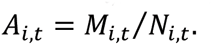

To calculate Ab gene expression, let *V* be the total number of unique genes that we detected belonging to a given Ab gene class (e.g., for IGHV, *V*=68; excluding alleles). For *v* ∈ [1, *V*], let *L_v_* be the length of gene *v*. If *m*_*v,i,t*_ is the total number of reads that map to Ab gene *v* with an e-value ≤ 10^−20^, then the expression level, *E*_*v,i,t*_, of Ab gene *v*, in patient *i* at time-point *t* was calculated as,

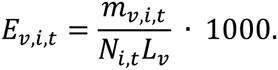

CDR3 abundance, or expression level, was calculated as the number of times a given CDR3 sequence was observed in patient *i* at time-point *t*, normalized by *N*_*i,t*_.

### Ab diversity

We used the CDR3 sequences in our dataset to estimate AbR diversity. However, there were different numbers of total reads sequenced for each patient/time-point, and diversity does not scale linearly with sample-size, so we could not simply normalize our diversity estimates by the total number of reads collected for that time-point, as was done for Ab abundance. To account for differing sample-sizes between patient/time-points we down-sampled our data until the number of reads for each patient/time-point was equal to the time-point with the least reads. We then calculated diversity from this down-sampled data. To account for possible stochastic effects of down-sampling, we analyze the mean of the 10 independently down-sampled diversity estimates.

We estimated overall AbR diversity by finding the mean pairwise genetic distance across the CDR3 sequence sets. Let *C*_*i,t*_ be the total number of unique CDR3 sequences found in patient *i* at time-point *t*, with *c* ∈ [1,*C*_*i,t*_]. Let *d*_*i,t,c*_ be the number of times the CDR3 sequence *c* was found in the given patient/time-point. 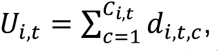, or the total number of CDR3 sequences detected. Additionally, let *s*_*i,t*_ be a list of actual CDR3 sequences. Antibody diversity, *D*_*i,t*_, for patient *i* at time-point *t* was estimated as,

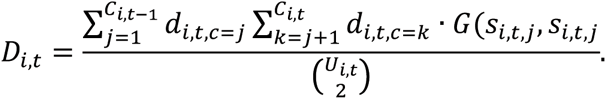

Where *G(x,y*) gives the genetic distance between the two CDR3 sequences, *x* and *y*. This was accomplished using the Needleman-Wunsch algorithm encoded by ‘needle’ in the EMBOSS package to first globally align sequences *x* and *y*. We then calculated ‘genetic distance’ by finding the percent of mismatches in this alignment, including gaps.

### Comparing B cell and PBMC CDR3 populations

We used a random sampling approach to test whether or not the CDR3 sequences from B cell and PBMC sample-types were random samples from the same population. Specifically, for a given patient we randomly choose a timepoint, then within this time-point, we randomly select one CDR3 sequence from the B cell dataset and one from the PBMC dataset, where the frequency of the CDR3 sequences determines the probability that it will be selected. We then calculated the genetic distance between these two sequences using *G*(*x, y*), as was done in the diversity calculation. This process was done 1,000 times to create a distribution of genetic distance values. To create null distributions we repeated this workflow, except sampled pairs of CDR3 sequences from the same population. We used the Mann-Whitney U test to determine if the B cell/PBMC distribution of genetic distances was significantly different from either of the nulls. This process was done for each of the patients.

### Test for identifying TIV-targeting Abs

The following method was used to identify both TIV-targeting V genes and TIV-targeting CDR3 sequences, so we shall henceforth use the notation ‘Ab-element’ to refer to either V gene or CDR3 sequence. For a detailed description of this FPCA based test see Wu and Wu [27], and associated R code [41]. Briefly, the test functions by first converting each of the Ab-element’s expression trajectories into a continuous function over the time-course, *t*, which we will call an ‘expression function’. This is accomplished by finding the linear combination of the naïve basis functions that best fit the observed expression data. These expression functions, *X(t*), can be expressed as,

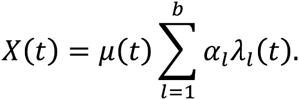

Where *μ*(*ί*) is a constant function that is equal to the mean Ab-element expression over the time-course, *a_x_* is the weight given to basis function *λ*_*ι*_(*t*), and *b* is the number of basis functions in the model.

FPCA is then performed on this set of expression functions, and the first set of eigenfunctions that explain at least 90% of the variance in the data were identified. Once this is done, *X’(t*) can then be re-expressed as a linear combination of this set of eigenfunctions that best fits the observed data.

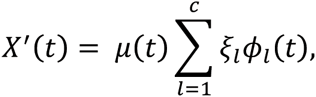

Where *ξ*_*ι*_ is the weight for each eigenfunction, *ϕ*_*ι*_(*t*), which is often referred to as the functional principal component score, and *c* is the number of eigenfunctions that form the set of eigenfunctions that together explain at least 90% of the variance in the data (such that their eigenvalues are non-increasing).

Once this is done, the task is then to determine if *X’*(*t*) is a better fit to the data than the null hypothesis. We define the null to be that the Ab-elements true expression function is *μ*(*ί*) (where the observed deviation around the mean is due to random error). Thus, the null hypothesis is *X*°(*t*) = *μ*(*ί*). We then determined which of the two hypotheses better fit the data by measuring the residual sum of squares (*RSS*) for the two models, *RSS’* and *RSS*°. The test statistic is given by,

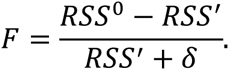

Where *δ* is a small constant that is meant to stabilize the variance of *F*, and is set to equal the variance of the Ab-elements observed expression values around its estimated expression function. Finally, in order to produce a null distribution of the test statistic, a permutation-based approach is used. The time-points are shuffled and this process is repeated. The Ab-elements whose *F* statistics were significant relative to the null distribution were deemed TIV-targeting. A Benjamini Hochberg correction for multiple tests was used on the p-values within a patient/gene class.

### Generation of literature-curated dataset of flu-targeting Abs

In order to characterize the diversity of Abs that have been reported to physically bind influenza, we scanned the literature and recorded the germline gene identity of all influenza-binding Abs that we found. As the generation of this literature-curated dataset qualifies as a meta-analysis, we created a separate Preferred Reporting Items for Systematic Reviews and Meta-Analyses (PRISMA) [42] statement that explicitly addresses each item in the PRISMA checklist in order to clearly outline the criteria used to select the studies that contributed to this meta-analysis. See the PRISMA statement and PRISMA Flow Diagram in the supplemental information for details on this meta-analysis.

### Test for a globally convergent V gene response

To determine if patients tend to use similar sets of genes to target TIV, we developed a statistic, which we refer to as ‘sum of gene significances’ (SGS), and is defined as the number of patients in which a given gene was found to be significant. Because we have 5 patients in our data, SGS is bound between 0 and 5. We computed the SGS value for each gene, and then compared the observed SGS distribution to its null. Generating a proper null model is paramount to any statistical test, and our task was to generate a null that takes into account the baseline frequencies at which the different V genes are expressed in a given patient, prior to vaccination. We chose to use a simulation-based null model, where we use day 0 gene frequencies to simulate artificial sets of TIV-targeting genes.

For our null simulations, we randomly sampled sets of genes for each patient, where the probability of sampling a given gene is equal to its frequency at day 0. The size of each null set of sampled V genes was the same as the number of genes that were found to be TIV-targeting in that patient. Thus, the null sets of V genes reflect underlying frequencies at which the genes were expressed *prior* to vaccination. This sampling process was repeated 1,000 times. We then combined these null sets of genes across patients and calculated the SGS statistic, establishing a null distribution of SGS. We then used a multinomial G-test to determine if the observed SGS distribution is significantly different than that of the null. If the significance threshold is exceeded by the observed data, then this suggests that the patients tend to use similar V genes when targeting TIV.

To generate the ‘naïve’ null distribution we treated each patient independently and then simulated SGS statistics under this model. Specifically, we estimated the probability that a V gene will be significant in a given patient by taking the fraction of V genes that were found to be significant in that individual out of the total number of V genes detected. We then simulated SGS statistics by walking through each patient and randomly assigning each patient as being significant based upon their estimated probability of V gene significance from the observed data. We then took the sum of these simulated significances to get the simulated SGS value. We did this 1,000 times and used the distribution of these simulated SGS values as our naïve null.

### Test for convergent response in individual V genes

This test is similar in spirit to the global test for V gene usage convergence (above), where the day 0 V gene usage is used to generate the null distribution. However, instead of a simulation based approach to generating this null distribution we develop a closed form solution. *P* is again the total number of patients in the study (5 in our case), and *p*_*£*_ is the relative proportion of a given V gene at day 0 in the tth patient (where *i* ∈ [1, *P*]). *S* is the set of identifiers for each patient, so *S* = {1,2,…, *P*], and *S_k_* is the set of all subsets of *S* that are of size *k*, so *S_k_* = {*x*|*x* ⊂ *s*, |*x*| = *k*], which represents all the different ways to choose *k* patients from *S*. If *X* is the random variable that describes the number of patients in which a given V gene is significant, then the probability of *X* under the null hypothesis is given by,

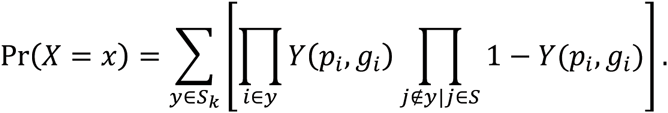

Where *Y(p_t_, g_t_*) gives the probability that a given gene will be found to be TIV-targeting in patient *i*, given that there where *g_t_* V genes that were found to be TIV-targeting in patient *i*. *Y*(*p_t_*,*g_t_*) is given as,

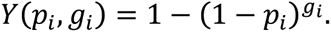

Essentially this can be thought of as a traditional urn problem in probability, where each patient is an urn that contains a given proportion of red balls. The probability of selecting a red ball from an urn is the probability of selecting a given V gene from a patient at day 0. The null distribution is modeled as follows: if *g_t_* is the number of draws made from each urn *i* (the number of TIV-targeting genes found for patient t), and p_£_ gives the probability of drawing a red ball from urn i (the relative frequency of the Ab gene in question at day 0), and *X* describes the number of urns from which red balls are drawn (the number of patients in which a particular V gene is identified as TIV-targeting); then the probability of *X* is the null distribution for SGS.

### Test for convergent CDR3 response

To test if two patients have sets of TIV-targeting CDR3s that are more similar to each other than would be expected by chance, we again utilized a methodology that hinges on sampling from the day 0 distribution. First, we calculate the mean pairwise genetic distance test statistic (commonly referred to as *π* in population genetics) between the two patients’ observed set of TIV-targeting CDR3s. If *X* is the set of TIV-targeting CDR3 sequences in patient x, and *Y* is that of patient *y*, then *π* between patient *x* and *y* was calculated as,

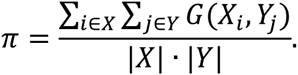

We then generate the null distribution for *π* by randomly sampling from the population of CDR3 sequences at day 0, where the frequency of each CDR3 sequence determines the probability that it will be sampled. The number of sequences that are sampled for each patient are equal to the number of CDR3 sequences that were found to be TIV-targeting for that patient. These sets of CDR3 sequences form a null set, and are solely informed by the baseline expression level of the CDR3 sequences prior to vaccination. We then calculate π between these two null sets, and repeat this sampling process 1,000 times to get a distribution of null π values. We can then assess significance of an observed π value between two patients by comparing it to their respective null distribution.

## Results

In this study, we implemented a pipeline to extract Ab sequences from RNA-seq data in order to take advantage of a unique densely sampled time-series dataset comprising RNA-seq data from PBMCs and sorted B cells of 5 patients vaccinated with the 2010 seasonal TIV over a time-course of 11 days [31] (Fig. S1). We use the high-resolution temporal information in these data in order to infer the elements of the AbR that are putatively targeting TIV. We then go on to test if the patients in this dataset exhibit more similar responses to TIV than would be expected by chance. That is, we test if these distinct AbRs exhibit convergence in response to the same vaccine.

### Quality control of bioinformatic pipeline

First, we validated that our bioinformatics pipeline (Fig. 1A, see methods for a detailed description) extracts meaningful AbR information from RNA-seq data. We hypothesized that the proportion of Ab encoding reads detected should correlate with the expected number of B cells in a given sample-type. We arbitrarily chose the day 7 time-point from patient 1, and applied our pipeline to the RNA-seq data from isolated B cells and PBMCs for this patient/time-point. As a negative control, we also applied our pipeline to RNA-seq data from human tissue cultured lung fibroblasts [43]. Our expectation was that the number of Ab sequences would decrease from B cells to PBMCs, and cultured lung fibroblasts would serve as a negative control with essentially no Ab sequences. Consistent with our expectation, we found that 1.25% of all reads from isolated B cells encode Ab (206,797 of 1.7×10^7^ total reads), PBMCs yielded 0.12% Ab encoding reads (16,214 of 1.4×10^7^ total reads), and cultured lung fibroblasts produced <0.001% Ab encoding reads (25 of 3.0×10^7^ total reads) (Fig. 1B).

### Broad AbR characteristics

We next sought to characterize how the AbR broadly behaves in response to TIV. To this end we measured Ab abundance (as the number of Ab mapped reads normalized by the number of non-Ab mapped reads, see methods) and Ab diversity (as mean pairwise CDR3 genetic distance, see methods) in each of the patients over the time-course (Fig. 2A, and 2B, respectively). We found that each patient had a characteristic peak in overall Ab expression around day 7, although the timing and severity of this peak varied dramatically among patients. Patient 3 had the most dramatic response, which had entirely subsided by day 7, while the response in patient 5 was much more gradual and less pronounced. We note that patient 3 was the only patient to have received the seasonal influenza vaccine for each of the prior 3 years and received the seasonal TIV as well as a monovalent vaccine the year prior to the collection date of the data used in this study. Together, these two vaccines had epitopes from two of the strains that were included in the TIV used in this study (Table S1). We note that the patients with the most dramatic increases in overall Ab abundance (patients 1 and 3), also tended to have a corresponding decrease in Ab diversity (Fig. 2B). We show in Figure 2C that this was largely due to clonal expansions of particular Ab lineages, presumably in response to the vaccine. Figure 2C also shows that the other patients seemed to have a predominantly polyclonal Ab response that did not substantially affect diversity. Though we cannot draw strong conclusions about the causes of a polyclonal or monoclonal response in this small sample size, future studies of larger cohorts could elucidate the causes behind this heterogeneity.

**Figure 2.**
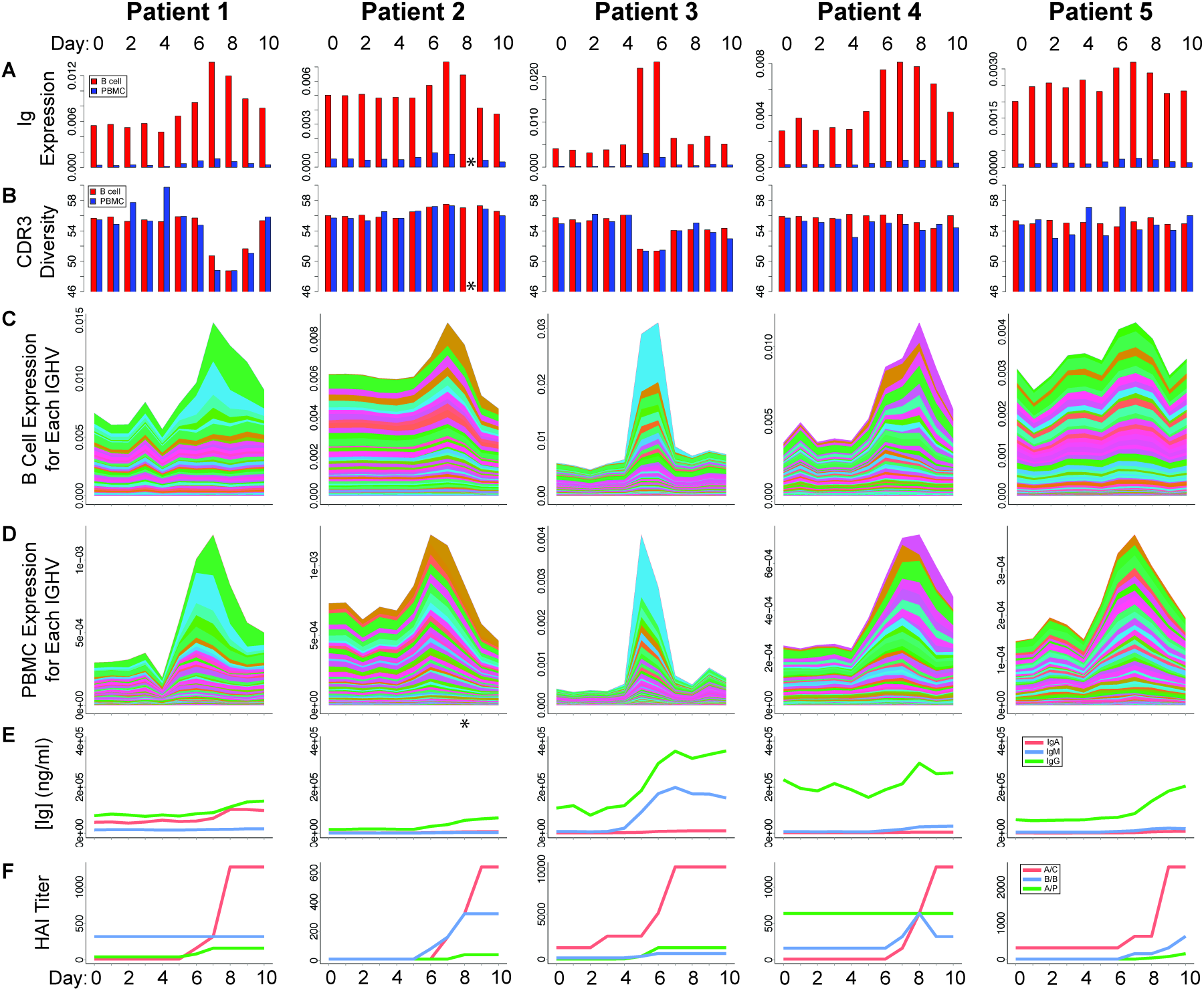
AbR response to TIV across patients. Different metrics were measured for each patient and at each time-point. Metrics are delineated by row, and patients are delineated by column. (A) Overall antibody abundance for each patient/time-point. (B) CDR3 diversity for each patient/time-point. B cells and PBMCs are shown in red and blue, respectively. (C and D) Stacked area charts showing the expression level for each IGHV gene for each patient/time-point. Colors, corresponding to IGHV genes, are comparable between patients and sample-types, and were sorted by absolute range (max - min). (C) B cell and (D) PBMC data. (E) ELISA results [31] giving the concentration of Abs that bind TIV for the A, M, and G Ab isotypes (red, blue, and green, respectively). (F) Hemagglutinin inhibition assay results [31] for the three different virus stains in the administered TIV, A/C: A/California/7/2009; B/B: B/Brisbane/60/2008; A/P: A/Perth/16/2009. *Patient 2 PBMC data at day 8 was unavailable due to sample processing error [31].

These results are consistent across both the B cell and the PBMC RNA-seq sample-types. Indeed, the Ab abundance and diversity levels for each of the patients and time-points are highly correlated between the two sample-types (Ab abundance: Kendall’s tau = 0.639, p = 8.715e-12, Ab diversity: Kendall’s tau = 0.366, p = 8.083e-05; Fig. S2), suggesting that the overall signal represents the underlying AbR diversity and expression patterns.

### Comparing B cell and PBMC CDR3 populations

Because B cells are a subset of PBMCs we can expect that the RNA-seq data from these two sample-types should yield similar Ab sequences. By checking to see if this is indeed the case in our data, we have another means to check the accuracy of our pipeline. In order to quantify the similarities between the B cell and PBMC sample-types, we focused on CDR3 sequence sets. Specifically, we statistically tested whether the CDR3 sequences from the B cell and PBMC datasets are drawn from the same population. We did this by finding the distribution of genetic distances between CDR3 sequences from different sample-types, and comparing this to the same distribution from CDR3s in the same sample type (see methods). We found that none of these three distributions are significantly different in any of the patients (p > 0.07, see Fig. S3). We thus conclude that PBMC and B cell datasets can reliably be used to extract Ab sequences from RNA-seq data.

### Clonal analysis

We next sought to analyze how each Ab gene is expressed over the time-course after vaccine administration. We calculated the expression level of each gene (as number of reads mapped to a given Ab gene normalized by the number of non-Ab mapped reads, see methods) in each of the patients, and at each time-point. We analyzed each class of Ab gene that could produce reliable alignments: V gene heavy chain (IGHV), V gene lambda light chain (IGLV), and V gene kappa light chain (IGKV). We were unable to detect an appreciable number of reads aligning to D or J genes with high confidence, which is likely due to their short lengths. We then generated stacked area charts to observe how the cumulative and individual V gene expression changes over time (Figs. 2C, and 2D for IGHV; Fig S4 for IGLV and IGKV). We find that the patients with the most dramatic Ab response (patients 1 and 3) also seem to have the largest clonal expansions in very few V genes, and that these expansions seem to explain a large portion of the overall increase in Ab abundance. This is particularly well illustrated in patient 1, where the peak in Ab expression is entirely explained by an increase in expression of 2-3 V genes. In addition, patient 1 shows a dramatic response in IGHV and IGLV (Fig. 2C, 2D, and S3A, S3B), but no response in the IGKV genes (Fig. S4C and S4D), suggesting that the handful of Ab lineages that responded to the vaccine in this individual (perhaps by chance) did not use the kappa light chain.

We performed an analogous analysis using CDR3 sequence data. We gathered all unique CDR3 sequences for each patient/time-point sample and calculated their expression level (see methods). We again generated stacked area charts to observe how the predominant CDR3 sequences change in expression over time (Fig S5). We found that these data largely recapitulate the gene expression data, where CDR3 expansions tend to occur around the same time as the increases in V gene expression. Patient 1 again shows a dramatic expansion in a single clonal lineage.

### Immunological assays

Given the robust signal in our clonal analysis, we sought to validate that the clonal expansions we observe in our data were indeed in response to TIV. Henn *et al.* [31] performed a variety of immunological assays using the sera from each patient/time-point sample. We used these data [44] to determine if the patients gain immunological reactivity against influenza around the same time as the clonal expansions occur in our data. Our analysis shows that vaccine-binding immunoglobin molecules tended to increase around the same time that the B cell clonal expansions (Fig. 2E). We next sought to establish that these clonal expansions conferred protectivity against influenza virus. Data from hemagglutinin inhibition (HAI) analyses using the three strains of influenza virus included in the TIV showed that protection to at least one of the strains was gained around the same time as the spike in clonal expansions (Fig. 2F). Together these data suggest that the V gene and CDR3 expansions we observe in our data are direct immunological responses to TIV.

### Identifying TIV targeting V genes

Given the robust signal that we saw in the V gene expression time-course data, we next established a methodology to systematically identify the V genes that appear to be targeting TIV. We utilized a method based on functional principal component analysis (FPCA), which was designed to identify differentially expressed genes over a time-course [27] (see methods for description). We found that it was often the case that the 1^st^ eigenfunction explained over 90% of the variance in the data (Fig. 3A and 3C). From this method we were able to identify the genes that seem to be most dramatically responding to TIV (Fig. 3B and 3D; Table 1). In almost all patients, the top genes identified in the B cell dataset (Fig. 3B) are replicated in the PBMC data set (Fig. 3D). We deem these to be TIV-targeting V genes. We then assessed whether or not these sets of TIV-targeting genes, or the AbR response in general, is more similar across patients then would be expected by chance.

**Figure 3.**
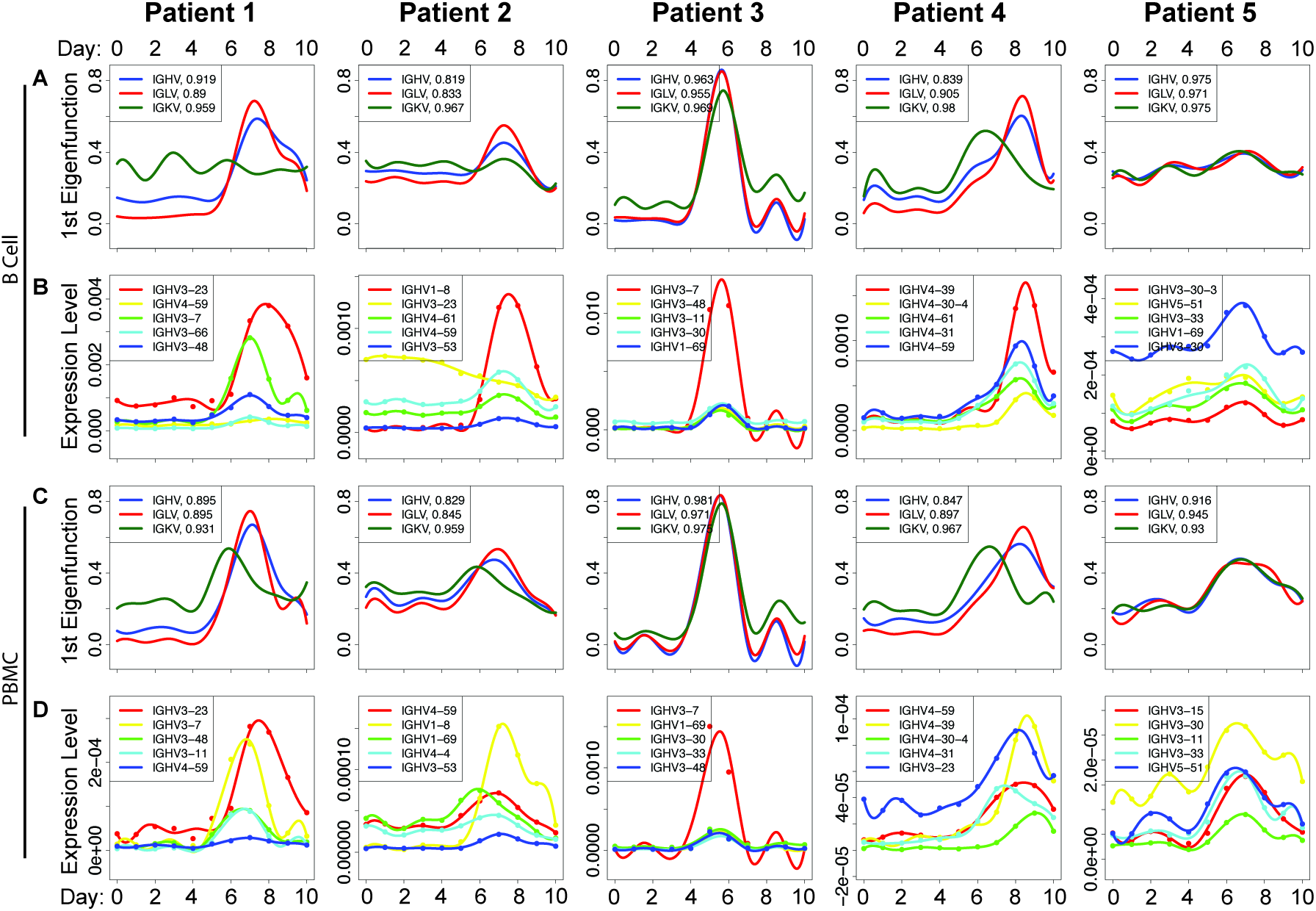
Identifying putative TIV-targeting V genes. (A) First eigenfunction in the B cell data for each patient and each V gene class. The proportion of the total variance explained by the first eigenfunction is listed in the legend after each respective class of V gene. (B) The top five scoring IGHV genes from the FPCA based test to identify TIV-targeting V genes; in the B cell data. The points show the observed data, and the solid lines show the best fitting gene expression function over time. V genes in legend are ordered by p-value, with lowest on top. P-values are based on a permutation test, see Wu and Wu [27] for details. (C and D) Same as (B and C) but from the PBMC data.

**Table 1.** Top 10 TIV-targeting heavy chain V genes.

| Gene Name | Lit Ab Freq. | Combined | Patient 1 | Patient 2 | Patient 3 | Patient 4 | Patient 5 |
| --- | --- | --- | --- | --- | --- | --- | --- |
| IGHV3-23 | 0.10345 | 1.08E-14 | 1.54E-05 | 4.55E-05 | 0.00529688 | 0.00010769 | 0.00015385 |
| IGHV1-69 | 0.23707 | 5.04E-14 | 0.00043284 | 0.00290909 | 0.00028125 | 6.15E-05 | 1.54E-05 |
| IGHV3-30 | 0.03879 | 7.19E-13 | 0.00153731 | 0.00154546 | 0.00025 | 0.00069231 | 1.54E-05 |
| IGHV3-7 | 0.04310 | 1.15E-12 | 0.00020896 | 0.00792424 | 1.54E-05 | 0.00010769 | 0.00389231 |
| IGHV4-61 | 0.00215 | 1.16E-12 | 0.00132836 | 7.58E-05 | 0.02834375 | 3.08E-05 | 0.00012308 |
| IGHV4-31 | 0.00431 | 1.78E-12 | 0.00110448 | 0.00424242 | 0.0015625 | 3.08E-05 | 7.69E-05 |
| IGHV3-48 | 0.01940 | 2.36E-12 | 0.00032836 | 0.00040909 | 0.0001875 | 0.00096923 | 0.00096923 |
| IGHV4-59 | 0.15302 | 2.77E-12 | 0.00011940 | 0.00013636 | 0.01929688 | 4.62E-05 | 0.00195385 |
| IGHV3-11 | 0.00431 | 3.14E-12 | 0.00037313 | 0.00074242 | 0.00025 | 0.00015385 | 0.00306154 |
| IGHV3-30-3 | 0 | 4.71E-11 | 0.00219403 | 0.01130303 | 0.00153125 | 0.00115385 | 1.54E-05 |

Lists the top ten scoring IGHV genes in the FPCA-based test for the B cell data. “Lit. Ab Freq.” lists the frequency for each of the V genes in the literature-curated dataset. “Combined” lists the p-values for each of the V genes after using Fisher’s method to combine the p-values from the FPCA-based test across all the patients. Patient 1-5 lists the p-values for each of the individual patients from the FPCA-based test. Genes are sorted by combined p-value.

### Testing for a convergent V gene response

Recently there have been several studies that have reported independent AbRs showing signals of sequence convergence when challenged with a similar antigenic stimulus [15,21,35,36]. This suggests that independent patients may use the same V genes to target similar antigens. Consistent with this, we found that V genes tend to have similar FPCA-based test scores across patients (Fig. S6). This suggests that the patients in our data are indeed using similar V genes to target TIV.

However, the sets of TIV-targeting V genes were statistically inferred using time-series data, and were not shown to physically bind TIV. To validate these findings, we searched the literature for Abs that have been experimentally shown to bind either an influenza vaccine or the influenza virus itself. Since most publications do not provide sequence information for the Abs they find, our analysis is limited to the germline genes from which the Abs originated. Our search resulted in 464 Abs that have been shown to bind influenza vaccine or influenza virus (Tables S2 and S3). We then compared the TIV-targeting V genes identified by our FPCA based test to the frequency of each V gene from our literature-curated dataset. Specifically, since each patient is approximately independent, we used Fisher’s method to combine FPCA-based p-values across patients. This results in a single p-value for each gene, where significance is increased if a gene is inferred to target TIV in multiple patients. Conversely, significance is diminished if a gene is heterogeneous across patients (Table S4).

We found that these combined p-values are positively correlated with IGHV gene frequency in the literature-curated dataset (Kendall’s Tau, B cell p = 3.115e-5, PBMC p = 2.502e-5). Moreover, we find that ∼60% of all Abs in our literature-curated dataset were composed of V genes that were inferred to be TIV-targeting in our analysis (Fig. 4A). In particular, we find that the genes IGHV1-69 and IGHV3-7, which have been shown to consistently target influenza epitopes in several independent studies [35,36,45–49] have the 2^nd^ and 4^th^ lowest p-values in the B cell data (Table 1), and 1^st^ and 2^nd^ lowest p-values in the PBMC data (Table S4), respectively. One of the publications that contributed to our literature-curated dataset used a combinatorial phage display library to select for influenza-targeting Abs (Throsby *et al.* [45]). This is different from the in-vivo selection process that occurs in humans, and thus could introduce unknown bias in the Abs from this study. We removed the data from this study and saw no qualitative difference in the outcome (Fig. S7). Together, these data show that (i) the V genes that we identify as TIV-targeting with our pipeline are consistent with previous findings in the literature, and (ii) that the patients from the Henn *et al.* dataset, as well as those from several other studies, tend to use similar V genes to target the influenza vaccine.

**Figure 4.**
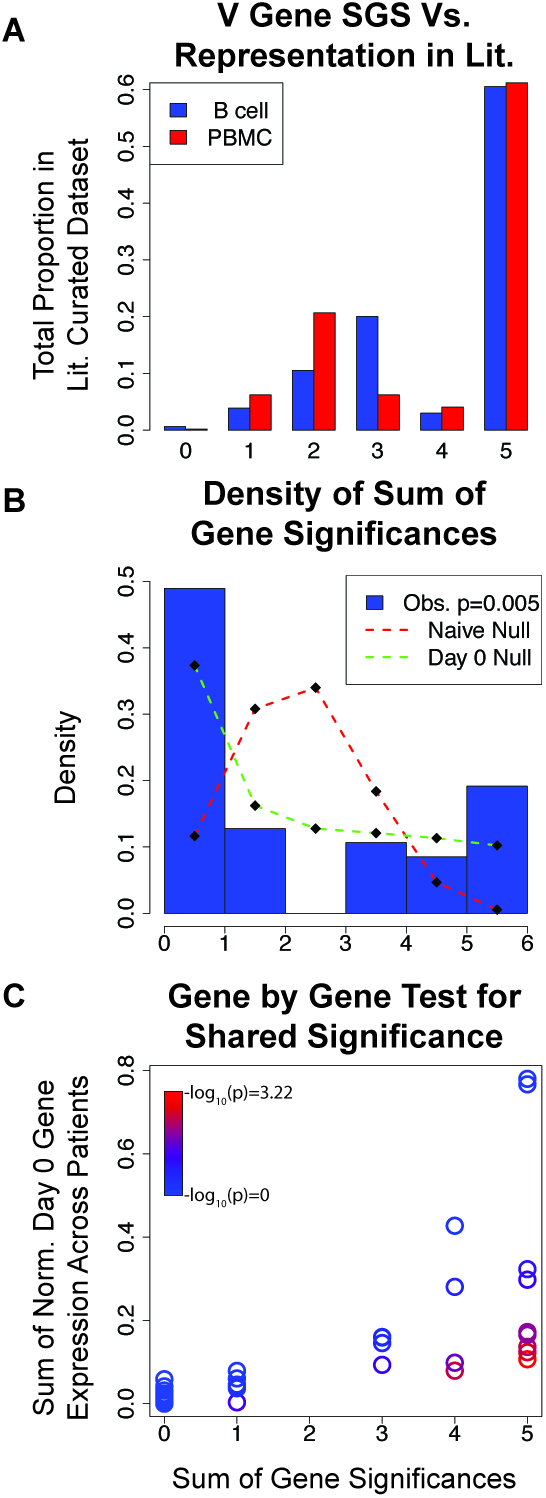
Identifying a convergent V gene usage signal across patients. The x-axis for all plots is the sum of gene significances statistic (SGS), which is the number of patients for which a given Ab gene was found to be significant. (A) Comparing our results for IGHV to the literature. For each SGS bin, this shows the proportion of the Abs in the literature-curated data that have V genes belonging to this bin. The genes that were significant in all 5 patients represented the largest proportion of the genes shown to be influenza binding in the literature. (B) Comparing observed SGS to the null distribution. Blue bars are a histogram showing the observed proportion of IGKV genes from the PBMC data belonging to each SGS bin. Red dashed line shows the ‘naïve’ null distribution of SGS if each patient were independent from one another (see methods). Green dashed line shows the null distribution of SGS if the baseline similarity in gene expression at day 0 is taken into account. The p-value in the legend shows the result of using a multinomial G test to compare the observed SGS distribution to that of the day 0 null. (C) Comparing the SGS value for each IGKV gene from the PBMC data to that of their respective null expectations. Color indicates the probability of the observed SGS under the null model (p-value, see methods for explanation of null model).

There are two reasonable explanations for this observation. The first is that some V genes have properties that make them naturally better at targeting TIV than others and are thus more likely to show a response across patients. The second is that patients tend to have similar V gene expression levels prior to vaccination, such that the Abs that are selected to respond to TIV tend to have similar V genes across patients simply due to this prior baseline similarity. We argue that this latter explanation has been underappreciated, and thus merits further scrutiny.

Suppose V gene expression levels are correlated across patients, independent of any antigenic stimulus. If Ab lineages were randomly selected to respond to an antigenic stimulus (the null expectation), then we would expect to see similar V genes responding to said antigenic stimulus across patients purely due to the underlying correlation of V gene expression prior to inoculation. We tested for correlations in V gene expression levels prior to vaccination (day 0), and found that they are highly correlated across patients (Fig. S8). We therefore developed a statistical test that will take into account the underlying similarity in V gene expression prior to vaccination when determining if the patients in our data tend to use more similar sets of V genes to target TIV than would be expected by chance (see methods). For each gene we find the number of patients in which the gene is found to be significant by our FPCA test (referred to as Sum of Gene Significances, or SGS, Tables S5 and S6). We then compare the observed SGS distribution to a null. We found that the observed SGS distribution was significantly different than the null for IGKV from the PBMC dataset (multinomial G-test, p=0.005; Fig. 4B; dashed green line vs. histogram) and was trending for IGHV in PBMCs (p=0.074), but we saw no evidence for convergent gene usage for IGLV (Fig. S9).

Given the mixed evidence for a global convergent signal in V gene response to TIV, we investigated each V gene individually (i.e., we test whether a given V gene was found to be TIV-targeting in more patients than expected by chance). Similar to our global V gene analysis, we used the gene frequencies at day 0 to construct our null distribution (the null was solved in closed-form, as opposed to simulating; see methods). We found that only two V genes showed a significant convergent signal after Bonferroni correction for multiple testing. These were IGHV3-66 on the heavy chain and IGKV3-NL1 on the kappa light chain, using the PBMC data (Tables S7 and S8). No lambda V genes showed a significant convergent signal with this test. In general, these significant V genes had a characteristic expression level trajectory where they started out as lowly expressed prior to vaccination, and then increased in expression postvaccination. This character of trajectory made it unlikely that the V genes would be selected to respond to the vaccine simply because they were abundant (or highly expressed) prior to vaccination, yet their increase in expression level after vaccination makes them likely candidates for responding to the vaccine. Both of these genes are absent from our literature-curated dataset. Interestingly, the V genes IGHV1-69 and IGHV3-7 — which have been reported in the past as showing a convergent signal when targeting TIV — are not significant in our test. This means that we cannot reject the possibility that these genes were found to be consistently targeting TIV simply due to their tendency to be highly expressed prior to vaccination.

Together, the results from our tests for a convergent signal in V gene usage show that some patients tend to use similar sets of V genes for particular gene classes, and that a couple of these V genes stand out. However, healthy skepticism is warranted for the interpretation of convergent signals in the AbR, as they are difficult to distinguish from baseline correlations that exist across patients.

### Testing for a convergent CDR3 response

We hypothesized that if the patients within this dataset are capable of using similar sets of V genes to target the same vaccine, then they may use similar sets of CDR3 sequences to target TIV as well. To answer this, we began by again using the FPCA-based test on our CDR3 expression data to identify the putative TIV-targeting CDR3 sequences. We were then left with a list of CDR3 sequences for each patient that appear to be targeting TIV. Our task was then to determine if these lists of TIV-targeting CDR3s were more similar between patients than would be expected by chance (see methods). Our results were mixed: patients 1 and 4 seem to have converged on similar CDR3 sequences to target TIV, whereas patients 2 and 3, and patients 1 and 3 seem to have diverged (Fig. 5 and S10).

**Figure 5.**
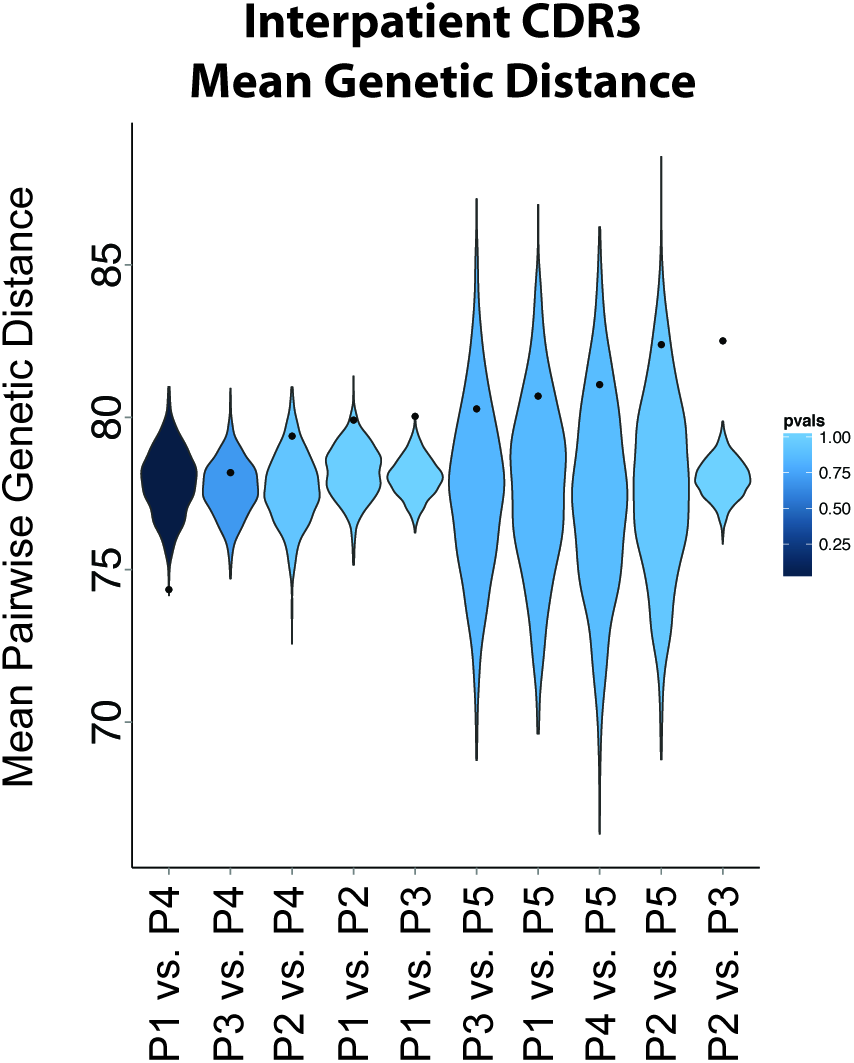
Testing for convergent CDR3 sequences across patients. Violin plots showing the null distribution of mean pairwise distance values for each pairwise patient comparison (see methods for how null distribution was created), for the B cell data. Black points indicate the observed mean genetic distance for the TIV-targeting CDR3 sequences between the patients compared. A point below the null distribution indicates convergent TIV-targeting CDR3 sequences, and above indicates divergent TIV-targeting CDR3 sequences. Patient comparisons are sorted by observed mean pairwise genetic distance. Distributions are colored by p-value with respect to the observed value. P1 vs. P4 p-value = 0.001.

## Discussion

We have mined and characterized the global AbR response to TIV in 5 individuals from RNA-seq data. We find that individuals exhibit a heterogeneous response to TIV. Some of the patients responded by expanding a few clonal lineages, while others responded with much more of a polyclonal character. Interestingly, patient 1, who had the largest clonal lineage expansion, was also the oldest patient (Table S1). This is in line with previous work showing that older humans tend to have larger clonal expansions in their AbRs [12]. While all the individuals’ overall Ab expression increased markedly post-vaccination, the timing and amplitude of this spike was variable. It is important to note that the patient with arguably the most dramatic response to TIV also had a relatively early spike in Ab expression, which had almost completely subsided by day 7. This is the time-point that immunologists typically collect samples for vaccine response studies (see Galson *et al.* [2] Table 1 for examples), and in this individual’s case the dramatic signal would have been all but lost if the traditional study design of pre-and post-vaccination time-points were used. This exemplifies the utility of study designs that emphasize dense, longitudinal sampling rather than cross-sectional sampling, as we would have missed much of the signal were there sparser sampling in the time-course. Further, as one decreases the number of time-points, it may become increasingly difficult to distinguish the signal from the noise, which would decrease the power to identify the elements responding to the stimulus.

While targeted sequencing of the Ab locus is unarguably the best way to illustrate the AbR, we, and others [33,34], have shown that a relatively simple bioinformatic pipeline can be implemented to characterize the AbR from RNA-seq data. This will hopefully provide investigators with the ability to leverage their RNA-seq data even further. Sequencing costs continue to plummet each year, however they still remain prohibitive for performing both targeted sequencing, and RNA-seq for the average project budget. If one were interested in overall, population level statistics of the AbR, such as abundance or diversity, or if one were interested in finding/observing the Abs that reach a high frequency in the AbR, we would argue that RNA-seq data is more than sufficient for these purposes. However, if one were interested in identifying rare Abs in the population, or needed full Ab sequences, then targeted sequencing of the Ab locus would be advised. In addition to prohibitive sequencing costs, targeted, deep-sequencing of the AbR remains a highly skilled method that involves a great deal of optimization, whereas RNA-seq has well vetted and broadly used protocols. In short, we hope that our method opens up the field of AbR analysis to a broader array of researchers.

The unique, densely sampled time-series dataset from Henn *et al.* [31] provided us with the ability to use functional data analysis methods to statistically identify putative TIV-targeting V genes. We found V genes that tended to be used among several patients in our dataset, and that these commonly used V genes are also prevalent in influenza targeting Abs collected from the literature. This finding suggests that we have identified V genes that indeed function to target TIV. This also raises the intriguing possibility that some V genes are inherently better than others at targeting TIV, as independent patients seem to be selecting the same V genes to target the vaccine. If this were true it would have interesting implications for the natural design and function of the diversity of genes in the AbR. For example, it could imply that instead of the different V genes providing the basis for an otherwise random exploration of sequence space when optimizing Abs, they could perhaps have evolved as ‘specialists’ for particular classes of antigens, such that when an Ab is comprised of a particular V gene it is pushed in a particular direction of antigenic space.

As interesting as a convergent signal may be, one must exercise great caution when searching for one. If correlations between individuals exist prior to the selection event, then these correlations must be controlled for in any convergence test. For example, consider a V gene that is highly expressed in many individuals prior to vaccination, and imagine that this V gene was found to be TIV-targeting in many patients. As we have pointed out, one does not know if the *reason* that this V gene was found to be TIV-targeting across patients is because it actually has a greater propensity to target TIV than other V genes, or because it was selected randomly due to its high prevalence in the individuals. It is certainly possible that the highly expressed V genes have a greater propensity to target TIV. Indeed, it is possible that the reason they are highly expressed is because of prior vaccinations/antigenic exposure. However, we argue that it is equally possible that some Ab genes have a high endogenous expression level independent of any antigenic stimulus. Because of this, we do not have the statistical ability to de-convolute these two possibilities. Increasing the number of patients in these types of studies would help ameliorate this problem. Alternatively, a synthetic AbR could be created that has a relatively even distribution of Ab elements, and tested for activity against TIV (or other antigens as well).

Our method of testing for a convergent signal in the AbR (i.e. using time-series data to identify the responding elements, then testing for convergence across patients) could be easily extendable to other systems. For example, this approach could be applied meta-genomic microbiome data in order to identify taxa that are consistently responding to some stimulus. It could also be applied to infections in order to see which sequence characteristics of a given pathogenic population are consistently responding to (or resisting) a drug.

## Conclusions

We have shown that AbR information can be harvested from RNA-seq data, that a densely sampled time-series can be used to identify the Ab elements that are responding to a stimulus, and that patients tend to use similar Ab elements to target the same vaccine, albeit in certain circumstances.

## List of abbreviations used

Ab: Antibody
AbR: Antibody repertoire
CDR3: Complementarity determining region 3
D: Diversity antibody gene segment
FDA: Functional data analysis
FPCA: Functional principal components analysis
HAI: Hemagglutinin inhibition assay
IGH: Heavy chain of antibody
IGHV: Variable gene on heavy chain
IGK: Kappa light chain of antibody
IGKV: Variable gene on kappa chain
IGL: Lambda light chain of antibody
IGLV: Variable gene on lambda chain
J: Joining antibody gene segment
mAb: Monoclonal antibody
NGS: Next generation sequencing
PBMC: Peripheral blood mononuclear cell
PRISMA: Preferred reporting items for systematic reviews and meta-analyses
RSS: Residual sum of squares
SGS: Sum of gene significances
SHM: Somatic hyper mutation
TIV: Tri-valent influenza vaccine
V: Variable antibody gene segment

## Competing interests

We (NBS and RDH) declare no competing interests.

## Acknowledgements

### Authors’ contributions

NBS wrote all scripts for analyses, developed/implemented all statistical tests, designed all figures/tables, and wrote this manuscript. The conception and design of this study, as well as all analyses herein are the result of a close collaboration between NBS and RDH. RDH helped draft this manuscript.

### Description of additional data files

The following additional data files are available with the online version of this manuscript. Additional data file 1 contains all supplemental figures and supplemental Tables S1 and S2 (Tables S3-8 are large so are instead included as separate CSV files), as well as the legends for all supplemental figures/tables that are cited in this manuscript. Additional data file 2 contains **Table S3; Literature-curated dataset of flu-targeting Abs.** Additional data file 3 contains **Table S4; P values for FPCA-based test.** Additional data file 4 contains **Table S5; TIV-targeting genes for each patient, B cell data.** Additional data file 5 contains **Table S6; TIV-targeting genes for each patient, PBMC data.** Additional data file 6 contains **Table S7; Results for individual gene test for convergence, B cell data.** Additional data file 7 contains **Table S8; Results for individual gene test for convergence, PBMC data.** The Tables in Additional data files 2-7 are formatted as CSV files, and the legends for these tables can be found in Additional data file 1. Additional data file 8 contains the PRISMA Statement that accompanies the meta-analysis portion of this study (see methods). Additional data file 9 contains the PRISMA flow diagram that accompanies the PRISMA Statement.

## Acknowledgments

This study was supported by grants from the National Institutes of Health (1R01HG007644 and 1R21HG007233), the Center For Aids Research (CFAR) at UCSF, and an Alfred P. Sloan Foundation Fellowship to RDH, as well as a Genentech Predoctoral Fellowship, and the Silvio Canonica Scholarship from the Swiss Benevolent Society to NBS. We kindly thank Dr. Manon Eckhardt for her valuable edits to this manuscript as well as her figure design contributions. We also offer warm thanks Dr. Lawrence Uricchio, and Raul Torres for their valuable input in the drafting of this manuscript. Lastly, we would like to thank the authors Henn et al. for publishing an invaluable dataset and making this study possible.

---

**Figure S1.**
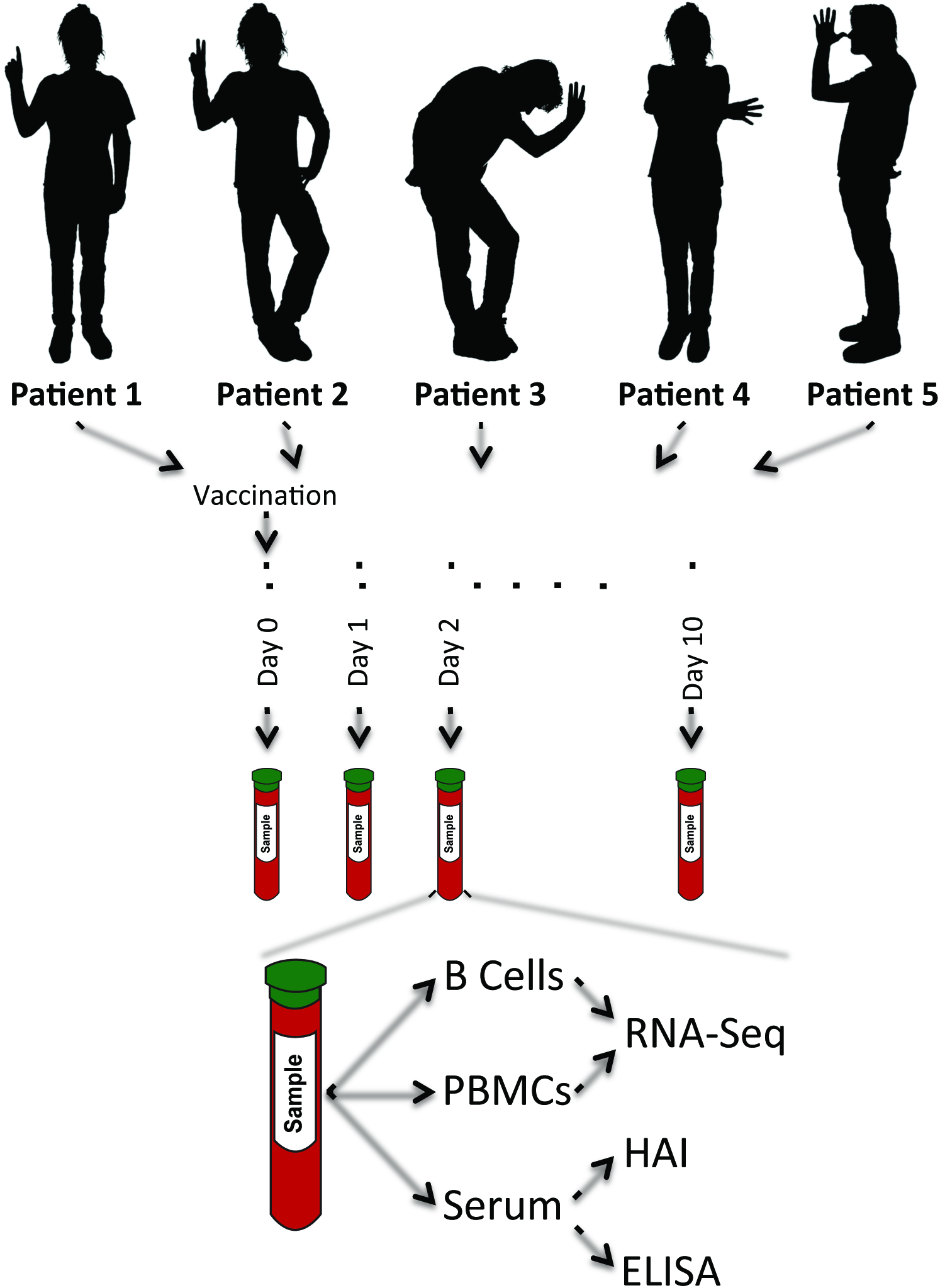
Study design. Schematic representation of the vaccination study. There are 5 patients and each patient is given the same vaccine. Whole blood is drawn immediately prior to vaccination, and each day for 10 days post vaccination (11 time-points total). Each patient/time-point sample is dived into three different sample types: Β cells, PBMCs, and serum. The Β cells and PBMCs are used for RNA-seq. The serum is used for immunological assays. *Note that patient avatars in no way reflect actual patients.

**Figure S2.**
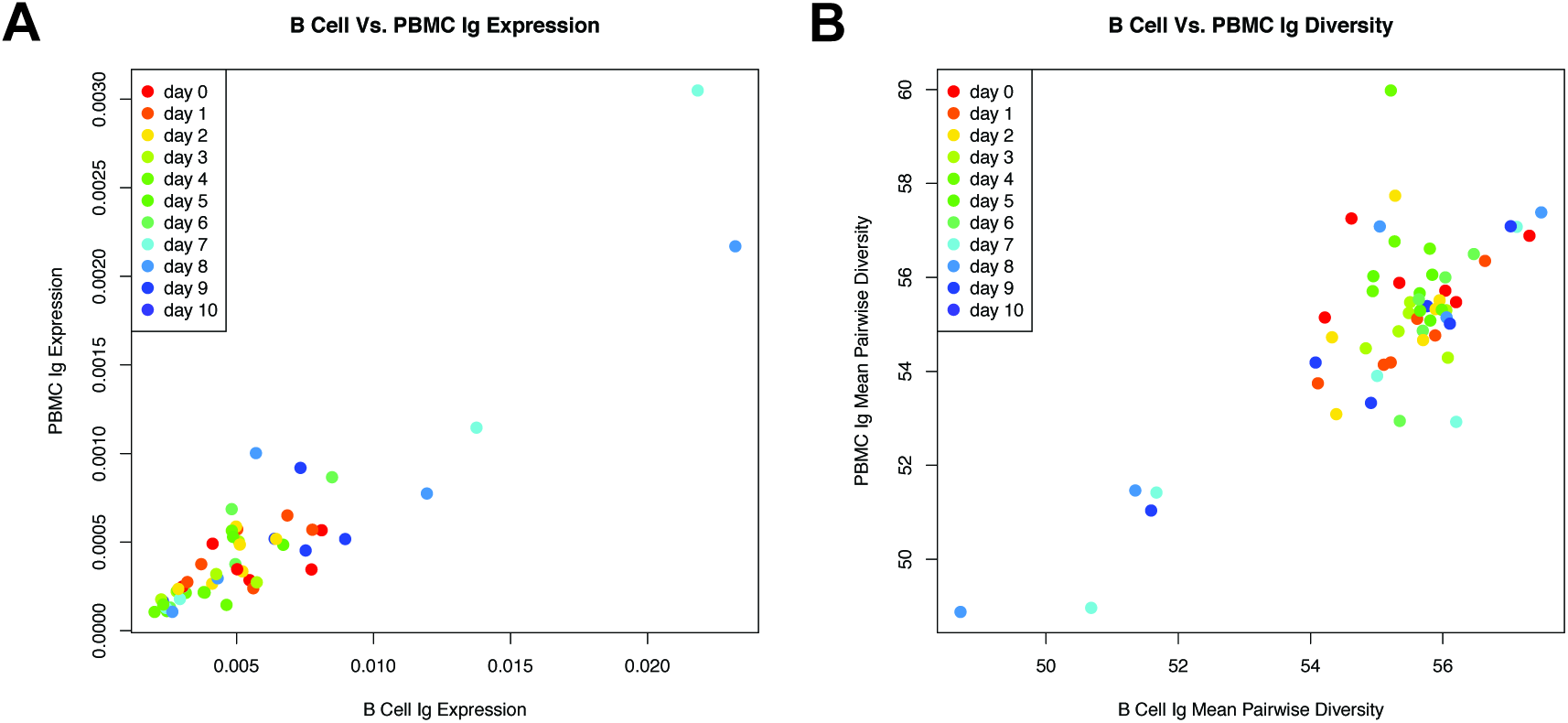
Correlation between B cell and PBMC AbR data. (A) Scatter plot showing the correlation between overall Ab expression in the B cell data to that of the PBMC data for each patient/time-point. Points are colored by time-point. (B) Same as (A) but showing correlation of mean pairwise diversity of CDR3 sequences between the B cell and PBMC data. See methods for how the diversity statistic was calculated.

**Figure S3.**
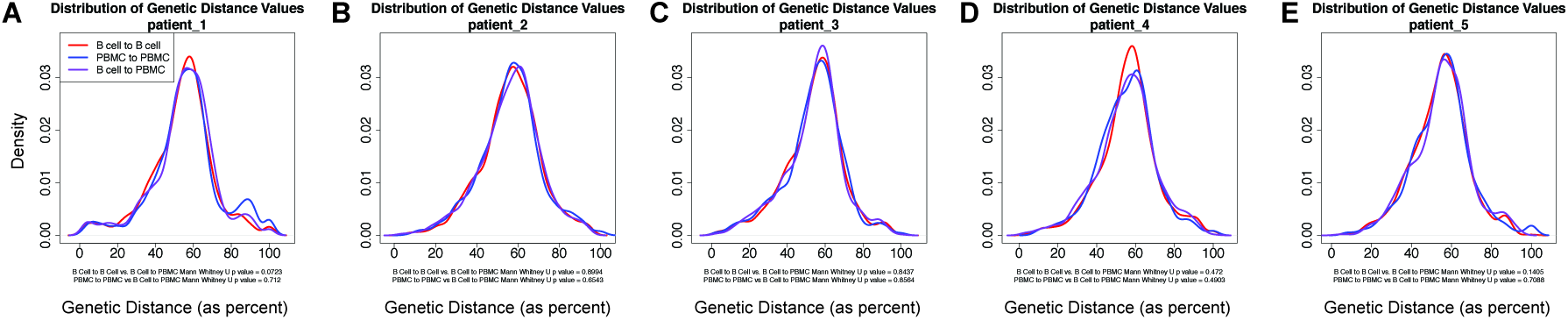
Comparing the B cell AbR to the PBMC AbR. Density plots showing the distribution of genetic distance values for randomly selected CDR3 sequences. Randomly selected sequences were matched by time-point. Red lines show the resulting distribution when comparing CDR3 sequences within the B cell data, blue lines show the same when using the PBMC data, and purple lines show the genetic distance distribution when comparing between datasets. Subtitle below the plots list the Mann Whitney U p values when comparing the red and the blue distributions to the purple. (A) Patient 1. (B) Patient 2. (C) Patient 3. (D) Patient 4. (E) Patient 5.

**Figure S4.**
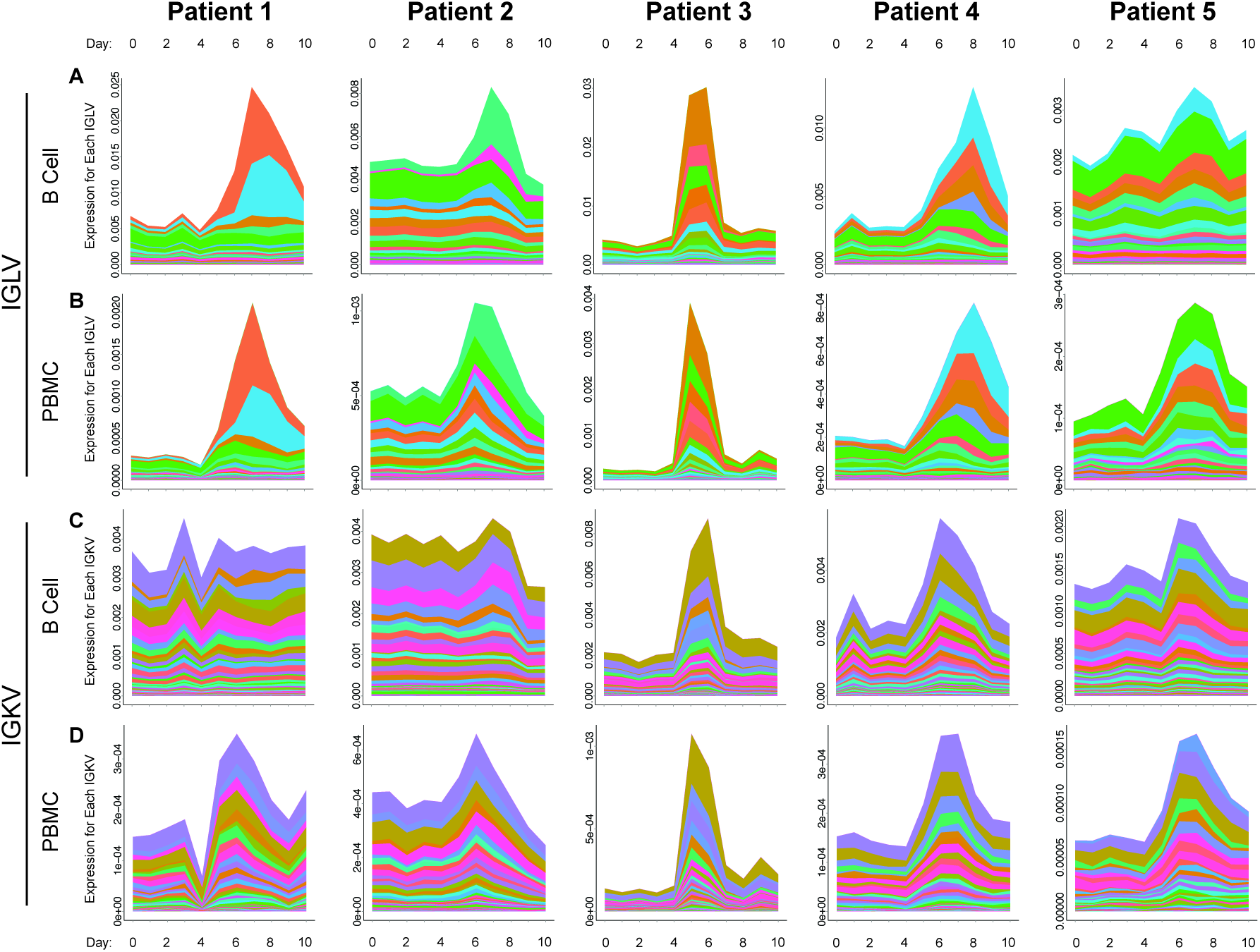
Lambda and kappa Ab gene’s expression over time. Stacked area charts showing the cumulative as well as individual Ab gene expression over time for both the lambda and kappa chains. (A and B) Stacked area charts for IGLV genes from the B cell and PBMC data, respectively. (C and D) Same a (A and B) but for the IGKV genes. All distinct colors for plots of IGLV genes correspond to the same genes (i.e. colors are comparable across patients and sample-types). The same is true for plots of IGKV genes.

**Figure S5.**
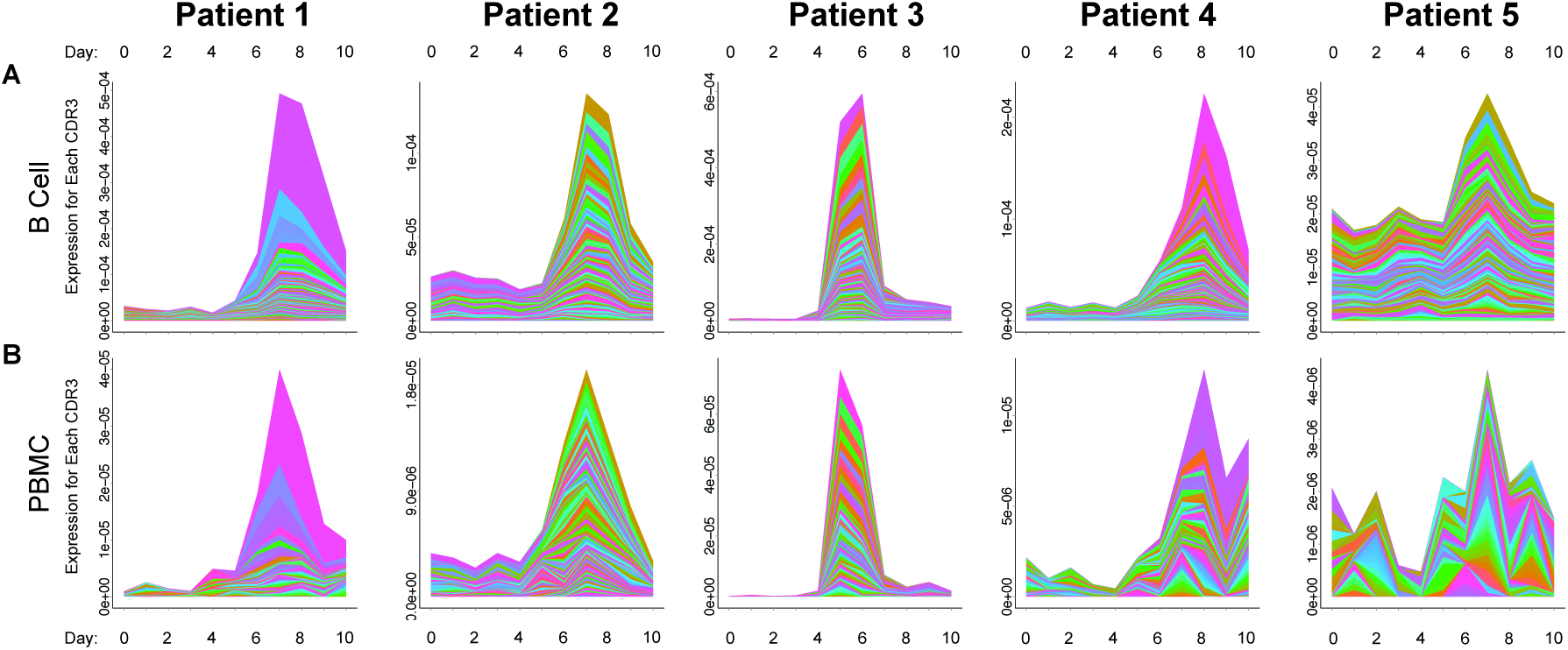
CDR3 expression over time. Stacked area charts showing the cumulative as well as individual expression level for the 100 most frequent CDR3 sequences in the data. (A) B cell data. (B) PBMC data.

**Figure S6.**
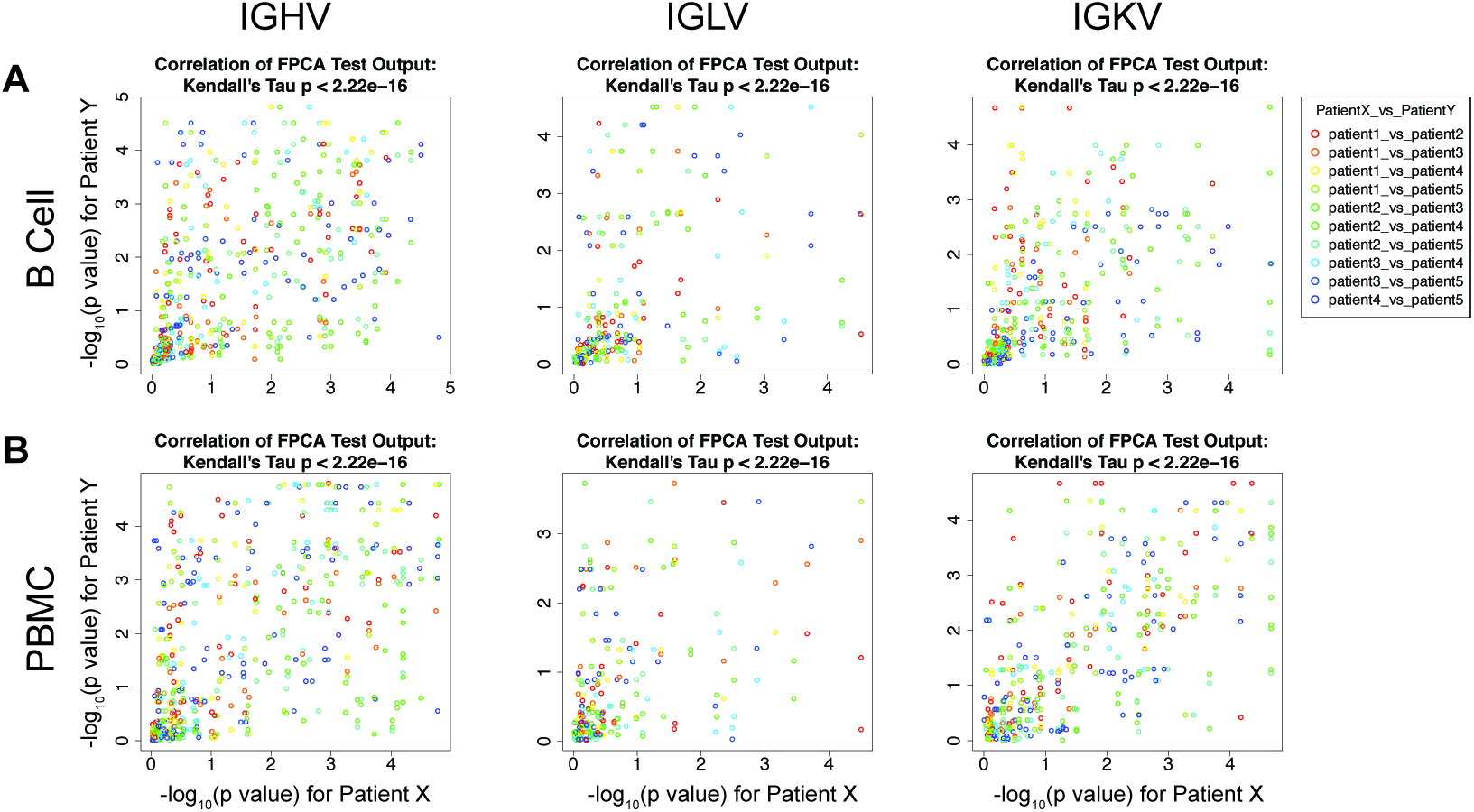
Correlation of TIV-targeting V genes across patients. Comparison of FPCA based test p values between all pairwise patients, for each V gene. For each pairwise patient comparison, these plots show the correlation of the p values from the FPCA based test for each of the genes. Points are colored by patient comparison. Correlation p value (Kendall’s Tau) is listed in the title of each plot. (A) Scatter plots for IGHV, IGKV, and IGLV genes’ p values for the B cell data. (B) Same as (A) but for the PBMC data.

**Figure S7.**
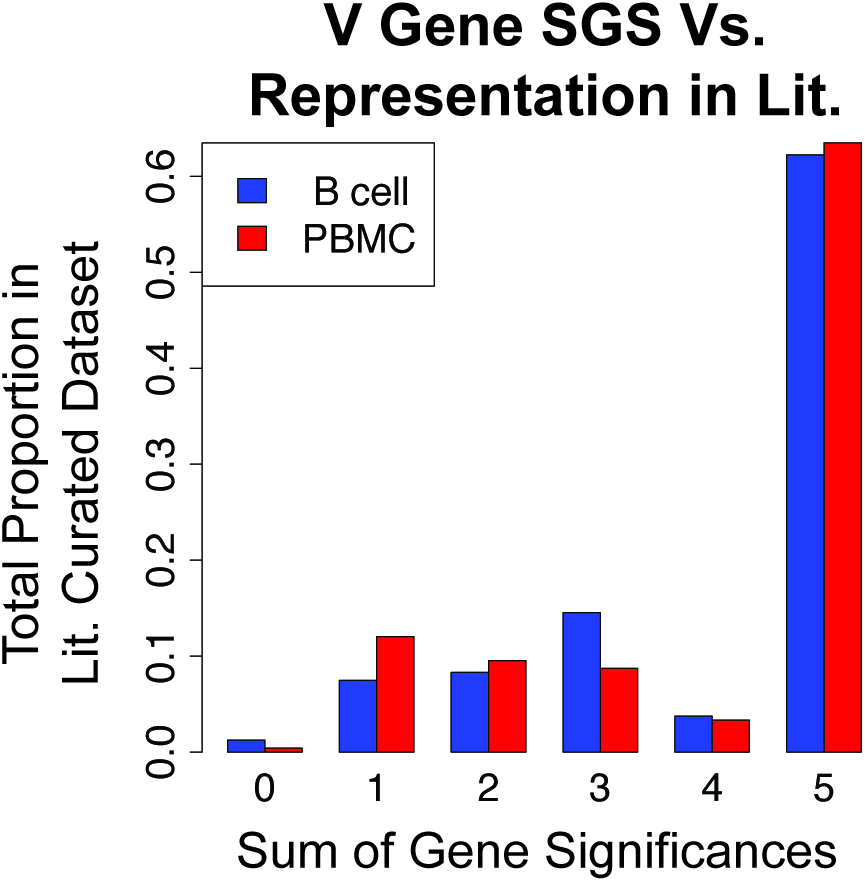
Comparison of SGS to literature-curated dataset, excluding Throsby *et al.* For each SGS bin, this shows the proportion of the Abs in the literature-curated data that have V genes belonging to this bin. The genes that were significant in all 5 patients represented the largest proportion of the genes shown to be influenza binding in the literature.

**Figure S8.**
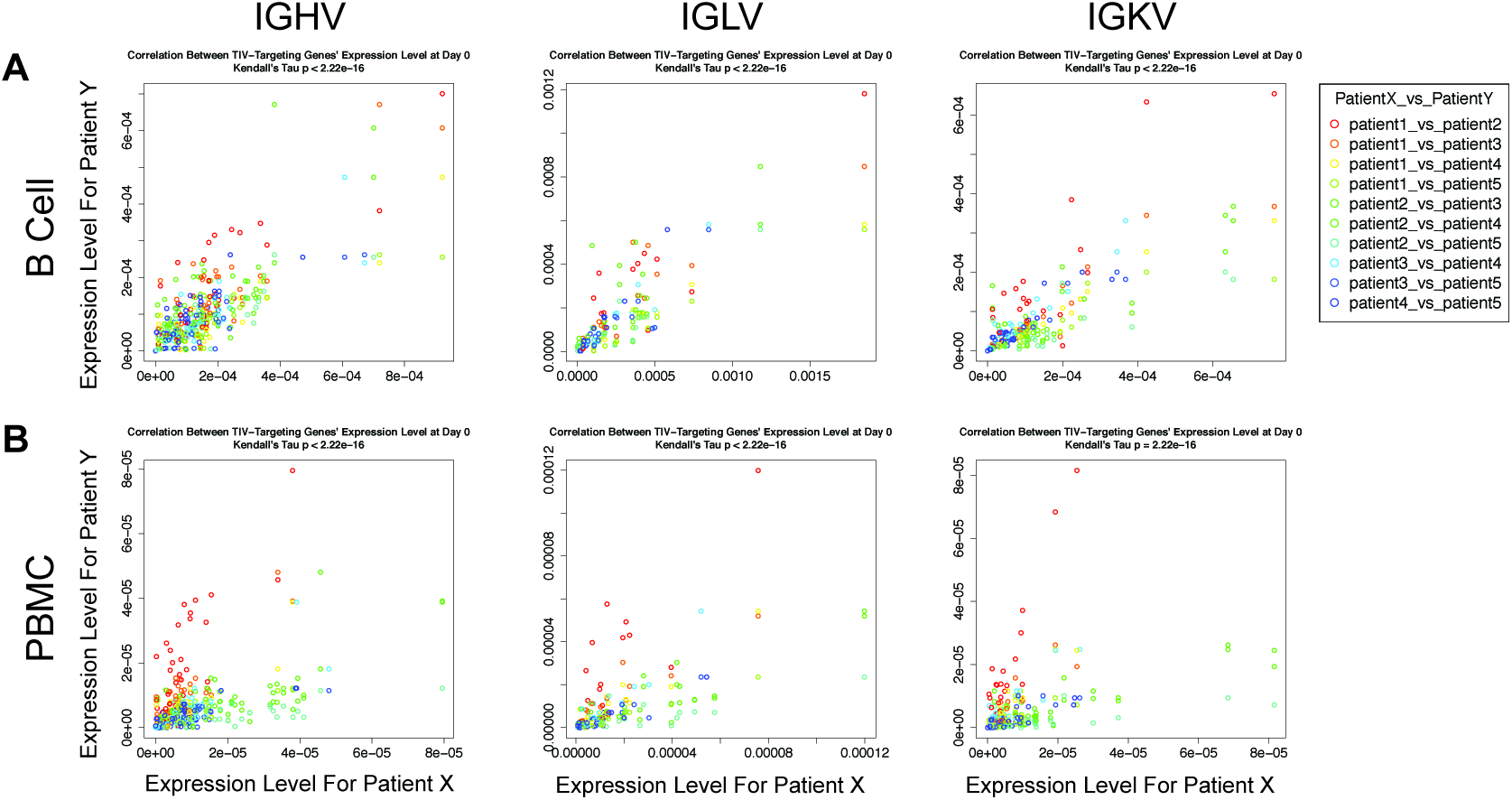
Baseline correlation of Ab gene expression at day 0. Comparison of day 0 expression level between all pairwise patients, for each TIV-targeting V gene. For each pairwise patient comparison, these plots show the correlation of the day 0 expression level for each of the TIV-targeting V genes. (A) Scatter plots for IGHV, IGKV, and IGLV TIV-targeting genes’ day 0 expression level for the B cell data. (B) Same as (A) but for the PBMC data.

**Figure S9.**
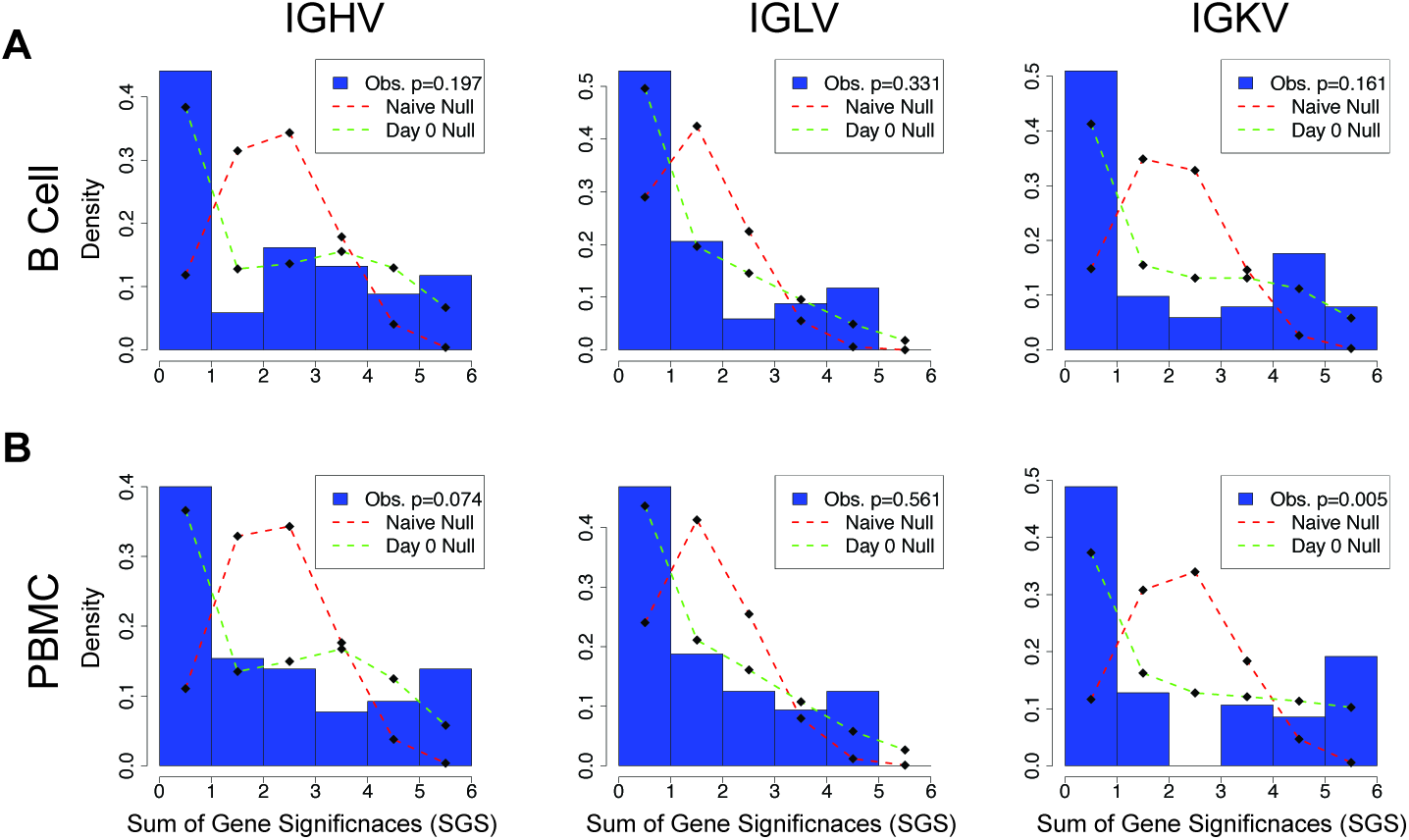
Testing for a global gene usage convergent signal. Comparing observed SGS to the null distribution. Blue bars are histograms showing the observed proportion of IGKV genes from the PBMC data belonging to each SGS bin. Red dashed lines show the null distribution of SGS if each patient were independent from one another. Green dashed lines show the null distribution of SGS if the baseline similarity in gene expression at day 0 is taken into account. The p values in the legends show the result of using a multinomial G test to compare the observed SGS distributions to that of the day 0 nulls. (A) Histograms of the IGHV, IGKV, and IGLV genes, for the B cell data. (B) Same as

**Figure S10.**
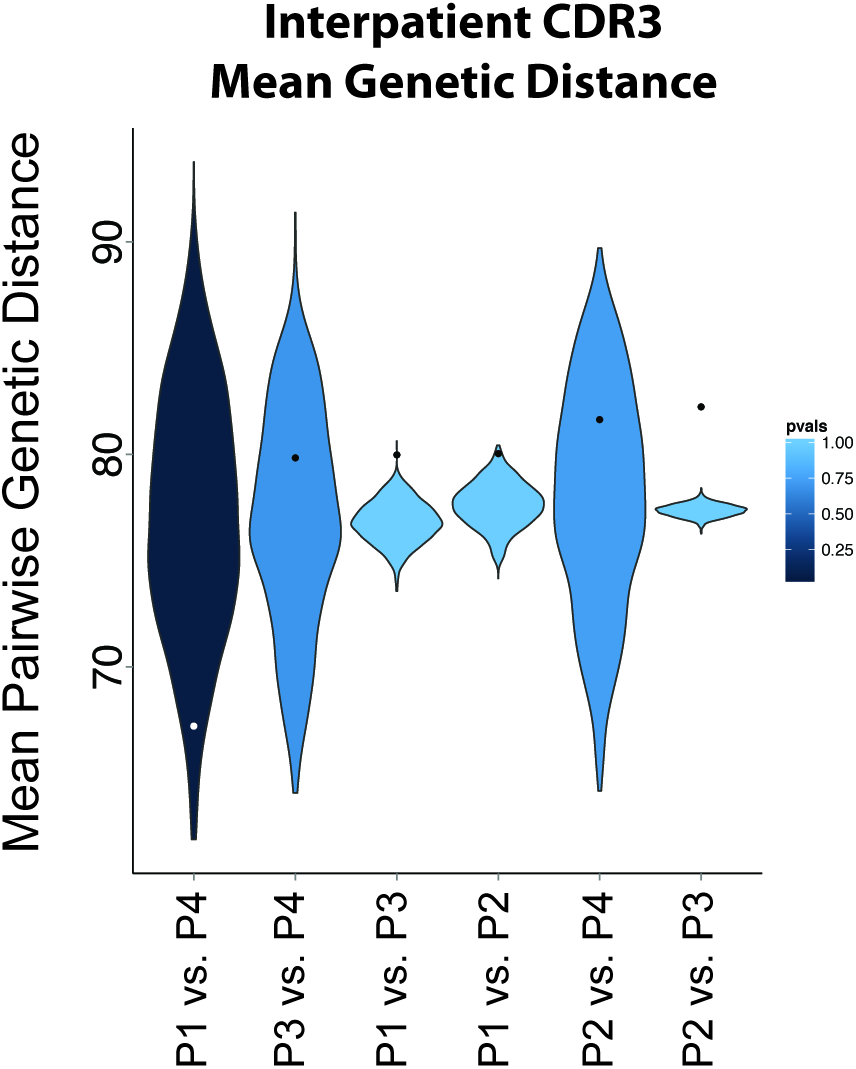
Testing for convergent CDR3 sequences across patients. Violin plots showing the null distribution of mean pairwise distance values for each pairwise patient comparison (see methods for how null distribution was created), for the PBMC data. Points indicate the observed mean genetic distance for the TIV-targeting CDR3 sequences between the patient comparison. A point below the null distribution indicates convergent TIV-targeting CDR3 sequences, and above indicates divergent TIV-targeting CDR3 sequences. Distributions are colored by p value with respect to the observed value. Patient 5 is absent because he had no statistically significant TIV-targeting CDR3 sequences in the PBMC data.

**Table S1.** Patient History.

| Subject vaccine history: |  |  |  |  |  |  |  |
| --- | --- | --- | --- | --- | --- | --- | --- |
| Subject ID | Age | Sex | Seasonal 2007 | Seasonal 2008 | Seasonal 2009 | Monovalent 2009 | Seasonal 2010 |
| Patient 1 | 47 | F |  | X | X |  |  |
| Patient 2 | 36 | F |  |  |  |  |  |
| Patient 3 | 20 | F | X | X | X | X |  |
| Patient 4 | 27 | M |  | X | X |  |  |
| Patient 5 | 23 | M |  |  |  |  |  |
| Strains used in vaccine: |  |  |  |  |  |  |  |
| A/H1N1 |  |  | A/Solomon Isl. /3/2006 | A/Brisbane/59 /2007 | A/Brisbane/59 /2007 | A/California/7 /2009 | A/California/7 /2009 |
| A/H3N2 |  |  | A/Wisconsin/67 /2005 | A/Brisbane/10 /2007 | A/Brisbane/10 /2007 |  | A/Victoria/210 /2009 |
| B |  |  | B/Malaysia/2506 /2004 | B/Florida/4 /2006 | B/Brisbane/60 /2008 |  | B/Brisbane/60 /2008 |

Top portion of table lists the demographics and vaccination history for each of the patients. If a patient has an ‘X’ under one of the vaccinations it means that that individual received the vaccine; a blank means that they did not. The lower portion of the table lists the strains in used in each of the vaccines. Colored entries mark the strains that were used in the study TIV.

**Table S2.** Contributions of each publication to the literature curated flu-

| Publication | IGHV | IGHD | IGHJ | IGLV | IGLJ | IGKV | IGKJ | Number Abs Retrieved |
| --- | --- | --- | --- | --- | --- | --- | --- | --- |
| Wrammert, 2011 | X | X | X | X | X | X | X | 46 |
| Krause, 2011 | X | X | X | X | X | X | X | 5 |
| Moody, 2011 | X |  | X | X | X | X | X | 160 |
| Corti, 2010 | X | X | X | X | X | X | X | 20 |
| Krause, 2012 | X | X | X | X | X | X | X | 5 |
| Yu, 2008 | X | X | X | X | X | X | X | 5 |
| Throsby, 2008 | X | X | X |  |  |  |  | 223 |

Lists all the publications that contributed to the literature-curated dataset. An ‘X’ under a given class of Ab genes means that that gene class was reported in the publication. If an entry is blank then the gene class was absent. “Number Abs Retrieved” lists the total number of monoclonal Abs that each publication contributed to the literature-curated dataset.

**Table S3. Literature-curated dataset of flu-targeting Abs.** Lists the germline gene identity for each of the Abs in the compiled literature-curated dataset. If an entry has an “NA” then that gene class was not reported in the publication. If an the genes for one of the light chains are left blank then that means that the given Ab was not composed of that light chain. The Wrammert, 2011 study did not amplify lambda genes, and thus none of the Abs from this study were composed of the lambda light chain.

**Table S4. P values for FPCA-based test.** Reports the p values for each of the detected V genes from the FPCA-based test to identify those that are TIV-targeting. “Lit. Ab Freq.” lists the frequency of each V gene in the literature curated dataset. “Combined All”, “Combined B cell”, and “Combined PBMC” list the p values when Fisher’s method is used to combine the p values across all patient/sample-types, across patients using B cell data, and across patients using PBMC data, respectively. B Cell Patient 1-5 lists the p values for each V gene for individual patients using the B cell data, and PBMC Patient 1-5 does the same using PBMC data.

**Table S5. TIV-targeting genes for each patient, B cell data.** Reports the TIV-targeting status for each of the V genes detected in our study, for each of the patients. If a given V gene has a 0 under a given patient, then that V gene was not deemed to be TIV-targeting by our FPCA-based test for that patient, whereas if there is a 1, then the V gene was TIV-targeting. SGS is the “Sum of Gene Significances” statistic, and is sum across each row. V genes are sorted by their SGS value.

**Table S6. TIV-targeting genes for each patient, PBMC data.** Same as Table S5, except using the PBMC data. See legend of Table S5 for description of data.

**Table S7. Results for individual gene test for convergence, B cell data.** Lists the p values for each V gene’s test for convergence, using the B cell data. “Obs. SGS” gives the observed SGS statistic for each gene. “Mode Null Dstrb. SGS” lists the mode of the calculated null distribution for the SGS statistic. “Sum of Day 0 Freqs.” gives the sum of the frequencies at day 0 across patients for a given gene. This is meant to provide a summary statistic to describe the level of expression (across patients) for each gene at day 0.

**Table S8. Results for individual gene test for convergence, PBMC data.** Same as Table S7, except using the PBMC data. See legend of Table S7 for description of data.

